# Structural and Dynamic Effects of PTEN C-terminal Tail Phosphorylation

**DOI:** 10.1101/2022.04.16.488508

**Authors:** Iris N. Smith, Jennifer E. Dawson, James Krieger, Stetson Thacker, Ivet Bahar, Charis Eng

## Abstract

The phosphatase and tensin homolog deleted on chromosome ten (*PTEN*) tumor suppressor gene encodes a tightly regulated dual-specificity phosphatase that serves as the master regulator of PI3K/AKT/mTOR signaling. The carboxy-terminal tail (CTT) is key to regulation and harbors multiple phosphorylation sites (Ser/Thr residues 380-385). CTT phosphorylation suppresses the phosphatase activity by inducing a stable, closed conformation. However, little is known about the mechanisms of phosphorylation-induced CTT-deactivation dynamics. Using explicit solvent microsecond molecular dynamics simulations, we show that CTT phosphorylation leads to a partially collapsed conformation, which alters the secondary structure of PTEN and induces long-range conformational rearrangements that encompass the active site. The active site rearrangements prevent localization of PTEN to the membrane, precluding lipid phosphatase activity. Notably, we have identified phosphorylation-induced allosteric coupling between the interdomain region and a hydrophobic site neighboring the active site in the phosphatase domain. Collectively, the results provide a mechanistic understanding of CTT phosphorylation dynamics and reveal potential druggable allosteric sites in a previously believed clinically undruggable protein.

## INTRODUCTION

Tumor suppressor phosphatase and tensin homologue (PTEN) is a tightly regulated dual-specificity (lipid and protein) phosphatase and keystone negative regulator of the critical PI3K/AKT/mTOR signaling pathway^1–3^. As a lipid phosphatase, PTEN catalyzes the dephosphorylation of phosphatidylinositol (3,4,5)-trisphosphate (PIP_3_) into phosphatidylinositol (4,5)-bisphosphate (PIP2), thereby regulating critical cellular functions and protecting against tumorigenesis^2,4^. Importantly, *PTEN* is frequently mutated in the soma and germline of individuals, where germline mutation carriers are predisposed to a difficult-to-recognize inherited cancer syndrome (Cowden syndrome) and neurodevelopmental disorder (NDD), such that *PTEN* is one of the most common genetic causes of autism^5–7^. The clinical heterogeneity associated with germline *PTEN* mutations is unified by the term *PTEN* hamartoma tumor syndrome (PHTS)^8,9^. PHTS is an autosomal dominant disorder variably characterized by macrocephaly, hamartomatous overgrowths, and malignant neoplasia, especially of the breast (85% lifetime risk in females), thyroid (35% lifetime risk), and endometrium (28% lifetime risk)^9–11^. Further, a risk of autism spectrum disorder (ASD) is significantly elevated in PHTS individuals, roughly 12-fold more common than general population risk^6,12^.

PTEN is a 403-amino acid composed of an N-terminal PIP_2_-binding domain (PBD, residues 1-15), a dual-specificity phosphatase domain (PD, residues 16-185), a C2 domain (C2D, residues 190-350), and a flexible carboxy-terminal tail (CTT) including the PDZ binding motif (PDZ, 401TKV403). The PD reveals the catalytic active site, which contains the WPD loop (residues 88-98), the highly conserved catalytic signature motif, HCxxGxxR (P loop, residues 123-130), and the TI loop (residues 160-171). Within the active site are three critical catalytic residues Asp92 (D92), Cys124 (C124), and Arg130 (R130). The C2D contains a short, unstructured intrinsically disordered region (IDR, residues 282-313). The CTT is also considered intrinsically disordered^13,14^, and contains two putative PEST domain sequences (residues 350-375 and 379-386), followed by the PDZ binding motif (**Figure 1**). Intrinsically disordered regions are dynamic, better described as an ensemble or cluster of conformations than as a single structure^15,16^. PTEN’s critical disordered regulatory regions are missing or incomplete in X-ray crystal structures^17–19^ (**Figure 1A**). Nevertheless, a multitude of biological functions and regulatory influences govern the many cellular processes of PTEN’s tumor suppression role, with CTT phosphorylation being a key regulator of catalytic activity^20–25^, stability^26^, and membrane binding^27–30^. Further, several studies reveal PTEN inactivation in carcinogenesis is regulated by CTT phosphorylation that alters protein-protein interactions (PPIs)^17,20–25^.

**FIGURE 1.**
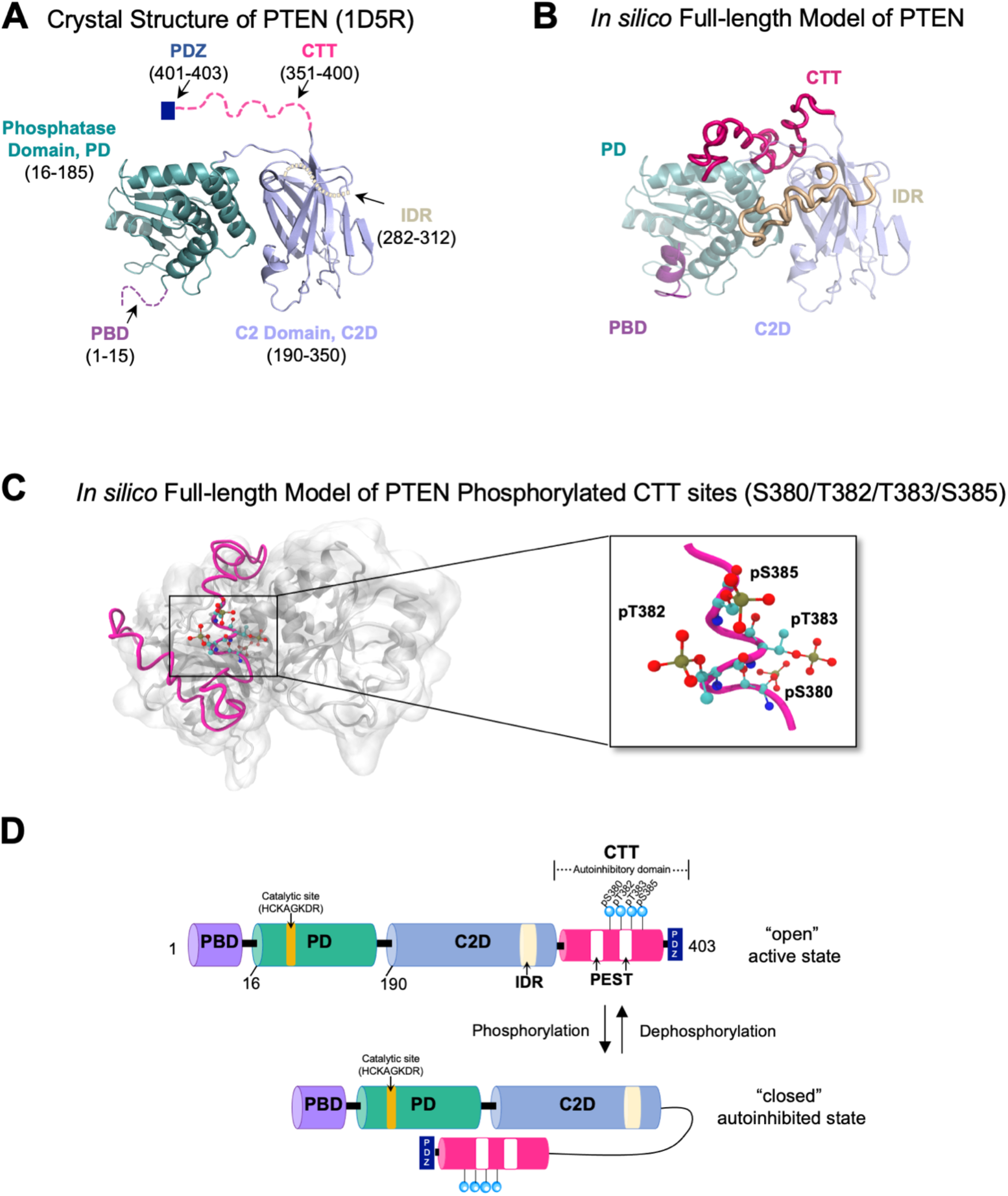
Structure and domains of PTEN. (**A**) The crystal structure of PTEN (PDBID: 1D5R) with an N-terminal catalytic phosphatase domain, PD (*cyan*, residues 16-185); a membrane-binding C2 domain, C2D (*light blue*, residues 190-350); and marked residues denoting missing regions. (**B**) *In silico* model of the full-length PTEN structure. The missing regions in PTEN, the PBD (*purple*, residues 1-15), the IDR (*tan*, residues 282-312), and the CTT (*magenta*, residues 351-403) were constructed using the I-TASSER program. (**C**) *In silico* fulllength model of PTEN with the phosphorylated Ser/Thr cluster in the CTT (*magenta*) indicated. Inset illustrates the four phosphorylated residues, pSer380, pThr383, pThr383, and pSer385. (**D**) Schematic illustration of PTEN in “open” and phosphorylation-induced “closed” autoinhibited

Biochemical and biophysical studies have revealed that the phosphorylation of a serine-threonine cluster (Ser380, Thr382, Thr383, and Ser385) on the PTEN CTT alters the protein conformational state from an “open” active state to a “closed” autoinhibited state in which the CTT folds over the active site and C2D (modeled in **Figure 1D**), resulting in reduced membrane localization and phosphatase activity^24,30–33^. This closed conformation appears to preclude productive CTT binding partner interactions, which further impairs enzymatic activity and disrupts downstream signaling pathways^14,34,35^. Yet very little is known about the mechanisms that govern phosphorylation-induced CTT conformational dynamics underlying such inactivation and no full-length high-resolution PTEN structure currently exists that shows the phospho-tail in the proposed conformationally closed state at atomic resolution. However, utilizing a combination of protein semi-synthesis, biochemical analysis, NMR, X-ray crystallography, and computation simulation, Dempsey *et al*.^18^, successfully crystallized and determined the X-ray structures of four semi-synthetic PTEN forms (PDBID: 7JUL, 7JUK, 7JVX, and 7JTX)^18^. Though their study offers tremendous insight into PTEN structure and provides a detailed framework on the structural impact of phospho-C-tail interaction, these structures only comprise residues 7-353 and lack the full-length PTEN structure. Moreover, they were unable to capture the phospho-tail and PTEN body (PD-C2) interactions by X-ray crystallography^25,36^.

Here, we employed molecular dynamics (MD) simulations on full-length *in silico* PTEN models (**Figure 1B and 1C**) in explicit water to determine mechanistically how phosphorylation at the CTT site (Ser/Thr residues 380-385) [**Figure 1C and 1D**] affects PTEN structural dynamics and ensemble. More specifically, we examined how the CTT conformational changes influence the active site loop dynamics. Towards these ends, we investigated three systems: (1) wild-type (WT) PTEN with unphosphorylated CTT, (2) PTEN with phosphorylated CTT (P-WT PTEN), and (3) phosphorylation-deficient PTEN that has its four phospho-sites mutated to Ala (PTEN 4A). Our *in silico* MD simulations support a structural model where the N-terminal PBD exists in an α-helical conformation, which agree with experimental results^18,37^, and also consistent with our integrative structural modeling^38^ and membrane MD simulation studies^39^. We show that CTT phosphorylation leads to a partially collapsed conformation, which alters the overall secondary structure of the PD and induces long-range conformational rearrangements that affect the active site. These active site rearrangements would prevent the localization of PTEN to the membrane and impede phosphatase activity. Notably, a phosphorylation-induced coupling is observed between the PD-C2D interdomain region and a region adjacent to the active site termed the “adjacent hydrophobic site” (AHS, framed by residues K13, R47, A126, G127, K128, T160, G165)^37^ within the N-terminal phosphatase domain. Molecular docking reveals that phosphorylation-induced conformational rearrangements do not prevent *in silico* association between the PTEN-PDZ motif and MAGI2-PDZ PTEN binding partner. However, close analysis of MD simulations of WT PTEN and P-WT PTEN reveals that phosphorylation reduces accessibility between PTEN and its PDZ. Our study sheds new light on the mechanistic interplay of CTT phosphorylation, conformational dynamics, and coupled underpinning of allosteric regulation in PTEN. Importantly, the detailed atomistic interactions and structural consequences of PTEN CTT phosphorylation identify druggable sites for binding allosteric modulators (or for allo-targeting), as a viable and currently unexplored potential treatment approach for individuals with PHTS.

## MATERIALS AND METHODS

### Preparation of the full-length *in silico* PTEN model

The full-length model of PTEN was generated previously by Jang and Smith *et al*.^39^. The procedure was as follows: the full sequence of the human PTEN (P60484, 403 residues) was obtained from UniProtKB/Swiss-Prot (www.uniprot.org/uniprot/)^40^. A PTEN X-ray crystal structure (PDBID: 1D5R)^17^ was utilized as a template to construct starting models of full-length WT PTEN and P-WT PTEN structures using the Schrödinger drug discovery software suite (Schrödinger, LLC, New York, NY, United States, 2017). The bound PIP_3_ analogue molecule in the crystal structure, tartrate (TLA), was removed and missing coordinates for residues 1-13 in the N-terminal PBD, residues 282-313 in IDR, and residues 352-403 in the CTT were modeled *in silico* by a hierarchal iterative templatebased threading approach using the I-TASSER webserver^41^ (**Figure 1B**). The CHARMM-GUI webserver^42,43^ was utilized to introduce phosphorylated CTT Ser/Thr cluster sites (S380/T382/T383/S385). The WT PTEN, P-WT PTEN, and PTEN 4A (S380A/T382A/T383A/S385A) systems were subsequently refined using Schrödinger Protein Preparation Wizard^44^ and Prime Wizard^45,46^ to assign bond order, add hydrogen atoms, and remove water molecules.

### Atomistic Molecular Dynamics Simulations

All-atom MD simulations were conducted using GROMACS 2020.2 simulation software^47^ with the CHARMM36m forcefield^48^ and a 2_fs timestep in explicit solvent. All systems were neutralized with NaCl ions inside a cubic box under periodic boundary conditions using the TIP3P solvent model. Each system was subjected to a stepwise energy minimization via the steepest descent method, and a series of four overall steps was performed as previously described^49^ to remove steric clashes and minimize forces. The minimized structures were subjected to a preequilibrium simulation for 350 ps with restrained backbones until the solvent reached 303 K. In the production runs, we employed the Nose-Hoover Langevin^50^ thermostat with a time constant of 1.0 ps to maintain the constant temperature at 303 K. The pressure was isotropically maintained at 1 atm using the Parrinello-Rahman barostat^51^ with a time constant of 5 ps and a compressibility of 4.5 × 10^-5^ bar^-1^. Lennard Jones potentials were smoothly shifted to zero from 1.0 to 1.2 nm and long-range electrostatic interactions were evaluated using the Particle Mesh Ewald^52^ method. For each WT PTEN, P-WT PTEN, and PTEN 4A structure production run, 1000 ns each (1 μs) of unrestrained NPT was performed at 303 K where bond lengths were constrained using linear constraint solver (LINCS)^53^ algorithm. Coordinates were saved every 20 ps, leading to 50,000 configurations for each WT PTEN, P-WT PTEN, and PTEN 4A system. Trajectory analyses were conducted using GROMACS utilities, VMD^54^, or PyMOL^55^ in-house scripts.

Each *in silico* PTEN system is composed of folded and disordered regions. The CHARMM36m forcefield^48^ used in our MD simulations is the closest to experimental measurement and demonstrate an accurate representation of intrinsically disordered proteins^56,57^. Intrinsically disordered proteins and regions have larger conformational ensembles available to them than folded proteins and are not readily represented by single structures^58,59^. The disordered PTEN regions are the PBD (residues 1-15), the IDR loop (residues 282-312), and the 50 amino acid CTT (residues 351-403), which are represented as an ensemble (**Figure S1**).

### Testing for production run reproducibility with different initial structures

The lowest energy structure model from I-TASSER was used as the initial structure for molecular dynamics simulation and subsequent analyses (**Table S1**). Duplicate production runs were performed to test reproducibility and structural stability assessed (**Table S2 and Figure S2**). A total of 6 μs simulations were performed, two simulations for each 1 μs system. To further sample the conformational space, the centroid structures from each most populated cluster ensemble from hierarchical cluster analyses served as initial structures of new MD simulations using the same protocol and repeated as (500 ns) replicas for WT PTEN, P-WT PTEN, and PTEN 4A (**Table S3**). Additionally, the second lowest energy structure from I-TASSER prediction and the lowest energy structure from a new full-length PTEN model using Rosetta FloppyTail^60^ application (**Table S4**, **see also Supporting Information**) served as initial structures of new MD simulations using the same protocol for 500 ns each. In total, 5 X 500 ns = 2.5 μs production MD simulations were performed as replicas for each system (three centroid structures and two independent models). The second lowest energy structure from I-TASSER and the lowest energy structure from Rosetta FloppyTail prediction have alternate ensemble CTT conformations (**Figures S3 and S4**) and were used to test the effects of different initial CTT conformations on the molecular dynamic simulations (**Supporting Information**). The simulations performed in this work are listed in **Tables S1-S4**. The conformational ensembles were assessed for all MD simulation runs (**Figures S1 and S4**).

### Elastic Network Model Analyses

The Elastic Network Model Analysis and subsequent analyses use initial WT PTEN, P-WT PTEN, and PTEN-4A structures based on the lowest energy I-TASSER structure. Both Gaussian (GNM) and Anisotropic (ANM) elastic network models (ENMs) identify the positions of network nodes with those of the Cα atoms and use uniform force constants of 1 kcal/mol Å^2^ for all the springs of the network. These models yield a spectrum of *N*-1 (GNM) or 3*N*-6 (ANM) nonzero modes of harmonic motions for a protein of N residues. Each mode *k* is characterized by a frequency and mode shape described by the respective eigenvalue, *λk*, and eigenvector, *μk*, of the Kirchhoff (GNM) or Hessian (ANM) matrices describing the network topology. The ANM is a coarse-grained ENM utilized to investigate protein dynamics and vibrational functional motions^61^. ANM calculations involved only Cα atoms and employed a uniform spring constant with a cut-off distance of 15 Å for interacting atoms, such that the overall potential of the system is a sum of harmonic potentials^62,63^. The centroid from the most populated cluster from hierarchical cluster analyses was utilized as input for the WT PTEN, P-WT PTEN, and PTEN 4A structure analyses. PRS was conducted using ANM with *ProDy*^64^. The PRS theory^65^ derives from Hooke’s law, applied to the ANM where F = HΔR or ΔR = H^-1^F. The displacement of all residues, ΔR^(*j*)^, was calculated in response to forces exerted on residue *i*, for all *i* = 1 to *N*. Allosteric PRS sensors and effectors were identified using the method described in recent work (General *et al*.^66^). PRS effectors and sensors were analyzed using *ProDy*^64^ using the first 20 slowest nonzero ANM modes^64^. ESSA^67^ was conducted using GNM with *ProDy*^64^ default parameters for inter-residue contact threshold distances (cut-off radii of 10 Å) between node pairs connected by elastic springs. An effective shift in global mode frequencies is defined for each residue as the mean 〈Δλ^(i)^_1-10_〉 over the softest ten modes (1 ≤ k ≤ 10), and a z-score (*z_i_*) is assigned to quantify the effect of ligand binding near residue *i* on global dynamics. *z_i_* = (〈Δλ^(i)^_1-10_) - *μ*)/σ where *μ* and σ denote the respective mean and the standard deviation of 〈Δλ^(i)^_1-10_〉 over all residues. First, the pockets of the specific structure are determined using Fpocket^68^ with its default parameters. Second, an ESSA score is assigned to each pocket, which corresponds to the median of the z-scores of the residues lining the pocket. Then the pockets are rank ordered based on the scores. As allosteric sites usually have relatively higher hydrophobic density, this Fpocket feature is used for further screening to determine a local hydrophobic density (LHD)^69^ z-score for each pocket based on all the pockets detected in the structure and then retain only those with positive z-scores. In the case of very close ESSA values, the LHD is used to refine the ranking further. As a result, all pockets are rank ordered in terms of their allosteric potential.

### Human PTEN PPI Interactome Analysis

Human PTEN PPI information was extracted from the BioGRID database v.3.5.174^70,71^. PTEN interactors with PDZ-binding domains (i.e. those interacting with the CTT domain of PTEN) were identified using PFAM annotations (http://pfam.xfam.org/)^72^, which were assessed by STRING analysis (https://string-db.org/)^73^ (**Table S5**). This list of PTEN-CTT interactors was cross-referenced with the literature and supplemented with two kinases that are known to regulate CTT phosphorylation, Casein Kinase 2 (CK2) and Glycogen Synthase Kinase 3-beta (GSK3β). The resulting list was again subject to STRING analysis to assess the interactions among proteins. The interactions were exported and subject to network analysis using Cytoscape 3.7.0 (https://cytoscape.org/)^74^. In the network, a node corresponds to a given protein and an un-weighted and un-directed edge corresponds to an interaction between those two proteins. PPI properties were assessed in terms of two common network topological centrality measures: *degree* and *betweenness centrality*. Degree centrality was defined as the number of interactions between interacting protein products with PTEN and is reflected by the size of the node (the size of a node is directly proportional to its degree). Betweenness centrality measures the importance of a node to the connection of different parts of a network – nodes with these characteristics are usually denoted as bottlenecks and considered hubs. The expansion in size and progression toward *red* color indicate increasing betweenness centrality.

### Molecular Docking and Construction of PTEN/PPI-PDZ Complexes

Molecular models of 1) WT PTEN/MAST2 PDZ, 2) WT PTEN/MAGI2 PDZ, and 3) P-WT PTEN/MAGI2 PDZ complexes were constructed using the full-length PTEN models (described above) and available NMR structures of PDZ domains. The PTEN/MAST2 complex (PDBID: 2KYL)^75^ comprised both a MAST2 PDZ domain (residues 1098-1193) and a PTEN CTT peptide (residues 391-403) and was utilized as a template to construct models. The PDZ-binding motif of PTEN was not in the freely available, ß-strand conformation needed to bind a PDZ domain. To overcome this difficulty, residues 1-390 of PTEN were fused using PyMOL^55^ to the ß-strand-like PTEN peptide from the PTEN/MAST2 complex (residues 391-403). This was done for both WT and P-WT PTEN models. The second PDZ domain of MAGI2 (PDBID: 1UEQ)^76^ was trimmed to residues 413-513 to remove disordered N- and C-terminal tails.

HADDOCK is a molecular docking program that can incorporate experimental data as restraints. Specifically, we used the HADDOCK 2.4 webserver^77^ to construct the three complex models. The initial guess for the binding interface residues were defined as “active” residues and treated as ambiguous restraints during the docking. The hydrogen bonds were defined as unambiguous restraints. For the WT PTEN/MAST2 model, the PDZ binding residues and hydrogen-bonds were based on the PTEN/MAST2 complex structure (PDBID: 2KYL) and defined as active residues and unambiguous distance constraints, respectively. For the WT PTEN and P-WT PTEN/MAGI2 models, the active and hydrogen-bonding residues of MAGI2 were defined by structural alignment of MAST2 and MAGI2 PDZ domains. The PTEN CTT’s (residues 351-403) were modeled as being fully flexible. 200 models were refined by HADDOCK in explicit water and the structures were clustered based on RMSD at the binding interface. The best structure for each model was chosen based on the HADDOCK score, cluster size (**Table 1**) and most reasonable strand-strand interaction between the PDZ and the PDZ motif. This structure was subsequently refined to minimize steric clashes and poor torsion angles using Rosetta3^78^. Quality of the structures were checked with MolProbity and post-refinement score^79^.

**Table 1.** Modeling and protein-protein docking quality assessment of MAST2 and MAGI2 PDZ for WT PTEN and P-WT PTEN systems.

| Complex | RMSD (Å) | Cluster Size | Buried surface area (Å<sup>2</sup>) | MolProbity Score |
| --- | --- | --- | --- | --- |
| WT PTEN/MAST2 PDZ | 2.2 (0.8) | 92 | 1218 (138) | 1.67 |
| WT PTEN/MAGI2 PDZ | 3.5 (1.2) | 97 | 1025 (95) | 1.71 |
| P-WT PTEN/MAGI2 PDZ | 2.6 (0.3) | 94 | 1115 (48) | 1.55 |

Standard deviations are given in parenthesis. RMSDs are based on the best four structures of the lowest energy/biggest cluster and the cluster size is out of 200 structures. Standard deviations are given in parenthesis.

### Structural Visualization and Analysis

VMD^54^ and PyMOL^55^ were used for simulation visualization. Graphs were generated in GRACE graphical plotting program (https://plasma-gate.weizmann.ac.il/Grace/)^80^.

## RESULTS and DISCUSSION

### CTT Phosphorylation Alters Structural Stability and Flexibility

Microsecond MD simulations were performed to elucidate the mechanistic effect of CTT phosphorylation on the structural dynamics of PTEN and uncover the underlying allosteric communication mechanism. We investigated three systems: (1) WT PTEN with unphosphorylated CTT, (2) P-WT PTEN, with phosphorylated CTT, and (3) a phosphorylation-deficient mutant of PTEN (PTEN 4A). While the WT PTEN explored a range of states in the vicinity of the starting structure, P-WT PTEN occupied a more “closed” state in line with the role of CTT phosphorylation in autoinhibition, and PTEN 4A occupied a more “open” state. The overall structural compactness was analyzed by calculating the radius of gyration (*R_g_*) [**Figure 2A**] where the WT PTEN system steadily fluctuates around an average of 23.1 Å, whereas the P-WT PTEN undergoes a gradual compaction reaching a lowest value of 22.2 Å around 700 ns, before relaxing to the same average of *R_g_* of about 23.1 Å, presumably after a major conformational change. We found that the *R_g_* of the entire protein for PTEN 4A initially increases to ~24 Å, but then undergoes a gradual compaction below both WT PTEN and P-PTEN. We also found that the highest *Rg* for the CTT for PTEN 4A and P-WT PTEN were found to be the same (~22.5 Å), whereas PTEN 4A has an overall higher *R_g_* compared to P-WT PTEN, which undergoes steady fluctuations before a sharp increase at 800 ns (**Figure 2B**), suggesting that the local structural arrangements of the CTT directly affects the overall protein conformation and dynamic flexibility. In contrast, the *R_g_* for the CTT in WT PTEN remains stable throughout the simulation after relaxing to an average of 17.5 Å around 200 ns.

**FIGURE 2.**
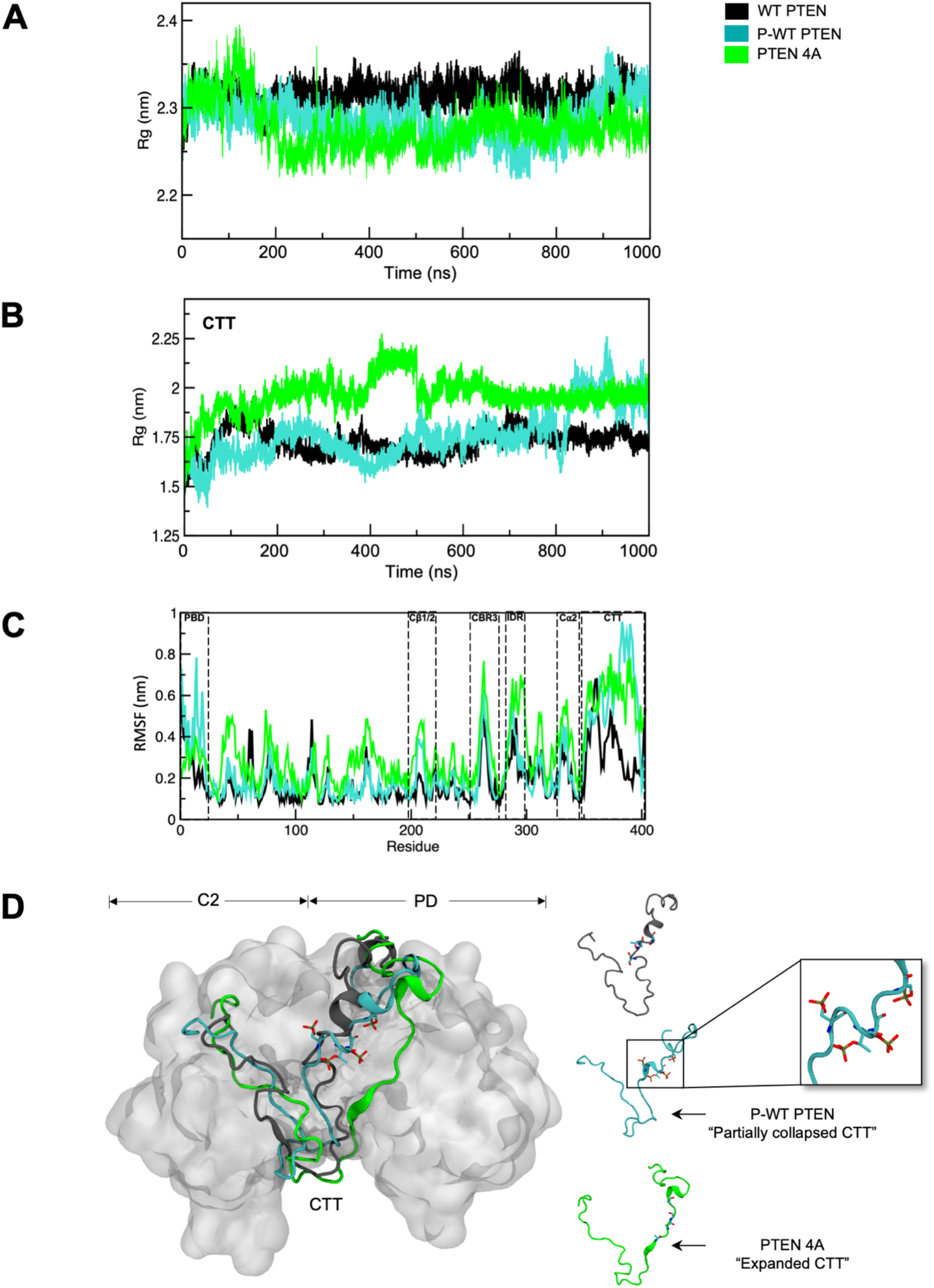
Conformational effects of PTEN CTT Phosphorylation. Time-evolution of (**A**) R*g* (entire protein), (**B**) R*g* (CTT only), (**C**) RMSF, and (**D**) Backbone superimposition of representative centroid structures of P-WT PTEN (*cyan*) and PTEN 4A (*green*) onto WT PTEN (*black*) reveals CTT conformational changes. Approximately 50,000 frames each representing a 2 fs time step were utilized for analysis.

The impact of CTT phosphorylation on the flexibility of PTEN was further explored via calculation of RMS fluctuations (RMSFs) of protein backbone atoms. The RMSF plots (**Figure 2C)** reveal that CTT phosphorylation affects the PBD, Cβ1/2, CBR3 loop (residues 260-269), several IDR residues, Cα2 loop (aka Motif 4, residues 321-334), and CTT regions for P-WT PTEN. Importantly, the Cβ1/2, Cα2, and CBR3 loops are critical to the membrane-binding interface in PTEN^81,82^, which suggests that high fluctuations in this region as a result of CTT phosphorylation may have some effect on membrane binding. Further, upon hierarchical cluster analyses, we superimposed the representative centroid conformations of each system to observe the structural changes as shown in **Figure 2D** and found major changes occurred in the CTT. The other structures of each cluster are shown in **Figure S1**, illustrating the variability within each conformational ensemble. We observed PTEN 4A to be highly flexible compared to WT PTEN and P-WT PTEN, specifically the PBD, pβ2-pα1 loop (aka R loop, residues 35-49), ATP-B motif (residues 60-73), several residues in the TI, Cβ1/2, IDR, Cα2, and CBR3 loops, as well as the CTT. We also found that CTT phosphorylation (P-WT PTEN) leads to a “partially collapsed” conformation of the CTT (**Figure 2D and Figure S1B**); whereas the phosphorylation-deficient CTT (PTEN 4A) leads to an “expanded” CTT and open/active conformer (**Figure 2D and Figure S1C**). Notably, we found the CTT structural ensembles for P-WT PTEN encompass a dynamic range of conformations, indicating a large diversity of backbone dihedral angle sampling (**Figure S1B**). A WT PTEN simulation starting from an alternate I-TASSER-derived initial CTT conformation (see **Materials and Methods, Supplemental Information**) produced clusters with similar compactness (root-mean-square deviation, RMSD) as the previous WT PTEN simulation (**Figure S4A and S4C**). Another alternate initial structure run was performed using a structure generated with Rosetta FloppyTail^60^ resulting in clusters with greater RMSD than the I-TASSER based simulations (**Figures S4B and S4C**), suggesting the possibility of a higher energy structure.

Next, we determined the simulation stability of each system (WT PTEN, P-WT PTEN, and PTEN 4A) and verified the time profiles of the RMSD of backbone atoms from the initial structure for each system. The structural dynamics of WT PTEN (*black*) and P-WT PTEN (*cyan*) are similar for a relatively long period (~700 ns) until P-WT PTEN displayed a sudden jump in RMSD at 750 ns and remains considerably higher thereafter, indicating that CTT phosphorylation has a major effect on the overall structural stability of PTEN (**Figure S1D**). The average backbone RMSD of WT PTEN remains around 4.4 (+/- 0.3) Å throughout the MD simulation relative to the initial structure, whereas that of the phosphorylated PTEN jumps from this value to about 6.0 (+/- 0.7) Å in the last 200 ns portion of the simulation. These results indicate that the P-WT PTEN ensemble has diverged from the initial structure more than the WT PTEN ensemble (**Figure S1A and S1B**). Similarly, the RMSD and *R_g_* is higher in the P-WT PTEN replica system compared to the WT PTEN replica system (**Figure S2A**). The PTEN 4A system has an overall higher RMSD compared to WT PTEN and P-WT PTEN (**Figures S1D and S2A**), suggesting that the CTT mutations at positions S380/T382/T383/S385 to alanine (“open/active” state) destabilize the overall PTEN structure. The CTT RMSD of each system reveals the P-WT PTEN displays a jump at 750 ns above both WT PTEN and PTEN 4A (**Figure S1E**), indicating the phosphorylation effects of the CTT govern the overall protein structural dynamics. However, the RMSD and *R_g_* are similar for a relatively long period until after 350 ns where the WT PTEN centroid replica system jumps, which indicates the overall structural stability changes as a result of a conformational change (**Figure S2B**).

### CTT Phosphorylation Promotes the Formation of a-Helices in the PBD and IDR

To further establish the effect of CTT phosphorylation on PTEN structure, we analyzed the time evolution of the secondary structures for the WT PTEN, P-WT PTEN, and PTEN 4A systems. To explore the structural variation of the CTT in the conformational ensembles, we performed clustering analysis of the respective trajectories. We found two dominant conformational ensembles holding >50% of total protein structures of each system. The percentage of the dominant conformations of WT PTEN, P-WT PTEN, and PTEN 4A, correspond to 95%, 51%, and 50%, respectively **(Figures 3A–C and S1A-C)**. The representative centroid structure from each conformational ensemble reveals the CTT in both the unphosphorylated WT PTEN and PTEN 4A is closer to the rest of the structure in comparison to P-WT PTEN, which is more outwardly displaced (**Figure 3A-C**). Further, the centroid structure reveals a “partially collapsed” CTT conformation in P-WT PTEN [**Figures 2D and S1B (*left* panel**)]. In contrast, the CTT of PTEN 4A is more “expanded”, which mediates long-range secondary structure changes in the PBD and IDR [**Figures 2D and S1C (*left* panel**)]. This agrees with previous experimental results that confirm the phospho-tail packs against the body of PTEN^25^. Further, our results support the simultaneous phosphorylation of the Ser/Thr cluster (S380/T382/T383/S385) described in Chen *et al*.^25^, which suggests that no single phospho-modification is dominant and that a tetra-phosphorylated WT PTEN could drive PTEN into a closed state.

**FIGURE 3.**
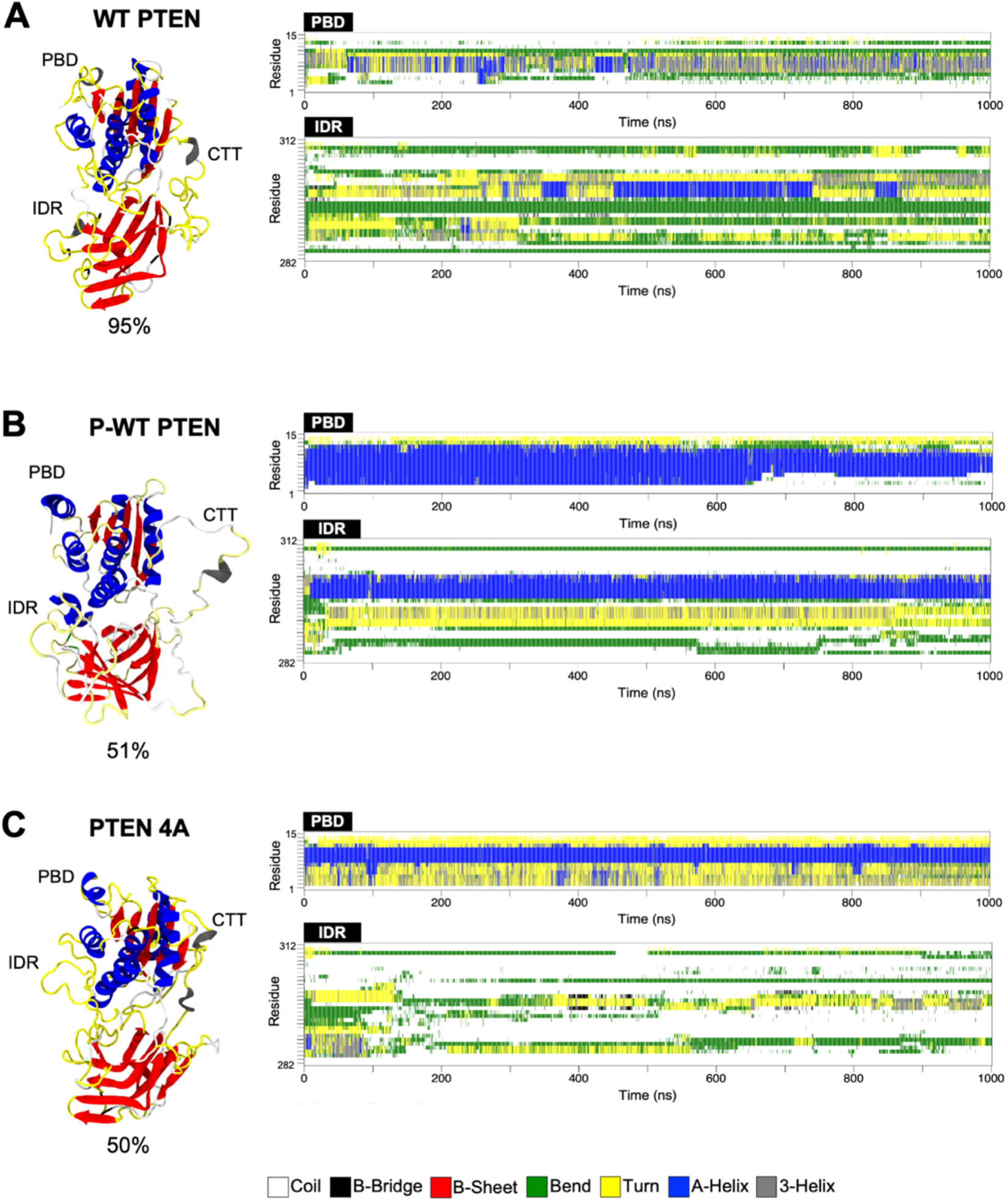
Secondary Structure analysis of (**A**) WT PTEN, (**B**) P-WT PTEN, and (**C**) PTEN 4A. CTT phosphorylation induces a-helical formation in P-WT PTEN. Secondary structure color-scheme was mapped onto the tertiary structure of the representative centroid structure for WT PTEN, P-WT PTEN, and PTEN 4A.

Of interest is the distinct increase in α-helical formation in the PBD and IDR regions in the centroid structure of P-WT PTEN compared to WT PTEN, which suggests CTT phosphorylation has a long-range stabilizing effect on PTEN structure (**Figure 3A and B; Figure S5A and S5B**). For the IDR, the P-WT PTEN system maintains a partial α-helical turn which persists after 200 ns compared to the WT PTEN (**Figure S5B**), suggesting CTT phosphorylation promotes IDR α-helical formation. Moreover, after 400 ns, the CTT reveals a 3_10_-helix proximal to the phosphorylated cluster site (Ser380, Thr382, Thr383, Ser385) that shifts into an α-helix at 800 ns (**Figure S6A**). For more detailed sampling, we utilized the centroid structures from each most populated cluster from hierarchical cluster analyses as starting structures of new MD simulations to further assess PBD and IDR structure, which agree with the initial structure results (**Figure S7**). Further, the secondary structure probabilities reveal that overall α-helical formation is increased in the PBD, IDR, and Cα2 loop (M4) regions for the P-WT PTEN system compared to WT PTEN as determined via DSSP secondary structure analyses (**Table 2 and Figure S8**). Throughout the simulations, the highest overall α-helical increase is seen in the PBD, which increased from 9% in the WT PTEN system to 63% in the P-WT PTEN system (**Table 2 and Figure 3A and 3B**). In contrast, a drastic loss in secondary structure is seen in the IDR region of PTEN 4A compared to both the WT PTEN and P-WT PTEN systems (**Table 2 and Figure 3C**). A modest increase in α-helical formation is seen in the PBD of PTEN 4A which persists throughout entire MD simulation yet increases from 9% in the WT PTEN system to 32% in the PTEN 4A system (**Table 2 and Figure 3A and 3C),** which suggests CTT mutation at positions S380/T382/T383/S385 to alanine destabilizes the overall structure of PTEN. Overall, our *in silico* MD simulation results reveal that phosphorylated CTT increases the presumed α-helical character in the N-terminal PBD and IDR regions of PTEN, which is in contrast to recent NMR studies^18^ where they demonstrate a phosphorylated CTT peptide diminishes the α-helical character of the N-terminus.

**Table 2.** Secondary structure probabilities for WT PTEN, P-WT PTEN, and PTEN 4A systems.

| System | Coil | β-sheet | Turn | α-helix | 3-helix |
| --- | --- | --- | --- | --- | --- |
| **WT PTEN** | 0.25 | 0.22 | 0.11 | 0.15 | 0.03 |
| PBD | 0.46 | / | **0.14** | **0.09** | 0.13 |
| IDR | **0.42** | / | 0.17 | **0.05** | 0.04 |
| CTT | **0.40** | / | 0.14 | 0.06 | 0.08 |
| WPD | 0.74 | / | 0.14 | / | / |
| P | 0.49 | / | 0.02 | / | / |
| TI | 0.66 | / | 0.05 | / | / |
| M1 | **0.34** | / | 0.01 | **0.52** | / |
| M2 | 0.96 | / | / | / | / |
| M3 | 0.96 | / | / | / | / |
| M4 (Cα2) | 0.34 | / | 0.33 | / | 0.05 |
| R | 0.69 | / | 0.25 | / | / |
| **P-WT PTEN** | 0.28 | 0.23 | 0.10 | 0.21 | 0.01 |
| PBD | 0.33 | / | **0.03** | **0.63** | / |
| IDR | **0.50** | / | 0.14 | **0.18** | 0.03 |
| CTT | **0.53** | / | 0.11 | 0.05 | 0.04 |
| WPD | 0.73 | / | 0.16 | / | / |
| P | 0.47 | / | 0.14 | / | / |
| TI | 0.64 | / | 0.18 | / | / |
| M1 | **0.17** | / | 0.01 | **0.83** |  |
| M2 | 0.95 | / | / | / | / |
| M3 | 0.95 | / | / | / | / |
| M4 (Cα2) | 0.33 | / | 0.17 | **0.17** | 0.16 |
| R | 0.65 | / | 0.13 | / | / |
| **PTEN 4A** | 0.28 | 0.24 | 0.11 | 0.19 | 0.02 |
| PBD | 0.17 | / | **0.39** | **0.32** | 0.10 |
| IDR | **0.71** | / | 0.09 | / | 0.02 |
| CTT | **0.59** | / | 0.07 | 0.03 | 0.06 |
| WPD | 0.75 | / | 0.16 | / | / |
| P | 0.49 | / | 0.25 | / | / |
| TI | 0.65 | / | 0.16 | / | / |
| M1 | **0.17** | / | / | **0.83** | / |
| M2 | 0.98 | / | / | / | / |
| M3 | 0.96 | / | / | / | / |
| M4 (Cα2) | 0.31 | / | 0.18 | **0.26** | 0.12 |
| R | 0.66 | / | 0.11 | / | / |

CTT phosphorylation and alanine mutations affect interactions between the pα3 helix (residues 102-115) and ATP-B binding motif (residues 60-73) in the PD (**Figure 4**). The pα3 helix is positioned directly behind the PTEN catalytic active site and participates in a rich hydrogen bond network with residues within the PD-C2D interdomain region^17^. Therefore, secondary structural changes that would alter the ATP-B binding motif and pα3 helix would disrupt the highly conserved rich hydrogen-bond network within the interdomain interface and alter PTEN catalytic function. An α-helix–coil transition occurs within the pα3 helix in both the WT PTEN and PTEN 4A systems, whereby the helix unravels into a disordered coil beginning at ~200 ns (**Figure 4A**). Conversely, CTT phosphorylation prohibits the unwinding of the pα3 helix in the P-WT PTEN system via a hydrogen-bond interaction between Tyr68 (Y68) of the ATP-B binding motif and Asp107 (D107) within the pα3 helix. The distance interaction between Y68(OH)-D107(Oδ1) was calculated throughout each simulation. We observed that the Y68(OH)-D107(Oδ1) bond maintained an interaction distance below 3 Å for a greater duration in WT PTEN and PTEN 4A versus P-WT PTEN (**Figure 4B**, *upper panel*). The hydrogen bond occupancy between Y68-D107 is higher in WT PTEN at 55% than in P-WT PTEN at 41% (**Table 3**). Further, the WT PTEN and PTEN 4A systems have a higher frequency of interaction compared to the P-WT PTEN system (**Figure 4C**). Additionally, an increase in α-helical formation is seen in the pα3 helix region for the P-WT PTEN system compared to WT PTEN (**Figure S9A and S9B, and S9C**). The rescue of the pα3 helix-coil transition appears to be governed by a sidechain interaction orientation between Y68-D107, where P-WT PTEN system predominantly prefers one orientation, and both the WT PTEN and PTEN 4A systems fluctuate between two orientations (**Figure 4B**). The pα3 helix is a highly conserved structural interdomain interface element that is crucial for maintaining protein stability and orientation of the catalytic active site^17,26^. In fact, this key interface consists of conserved residues frequently mutated in cancer, which indicates its critical role in PTEN function and the far-reaching consequences of mutations on the PTEN structure^17,27,83–85^. Further, site-directed studies have shown that alteration of the interactions between functional and highly conserved loops alter enzymatic activity^86^.

**FIGURE 4.**
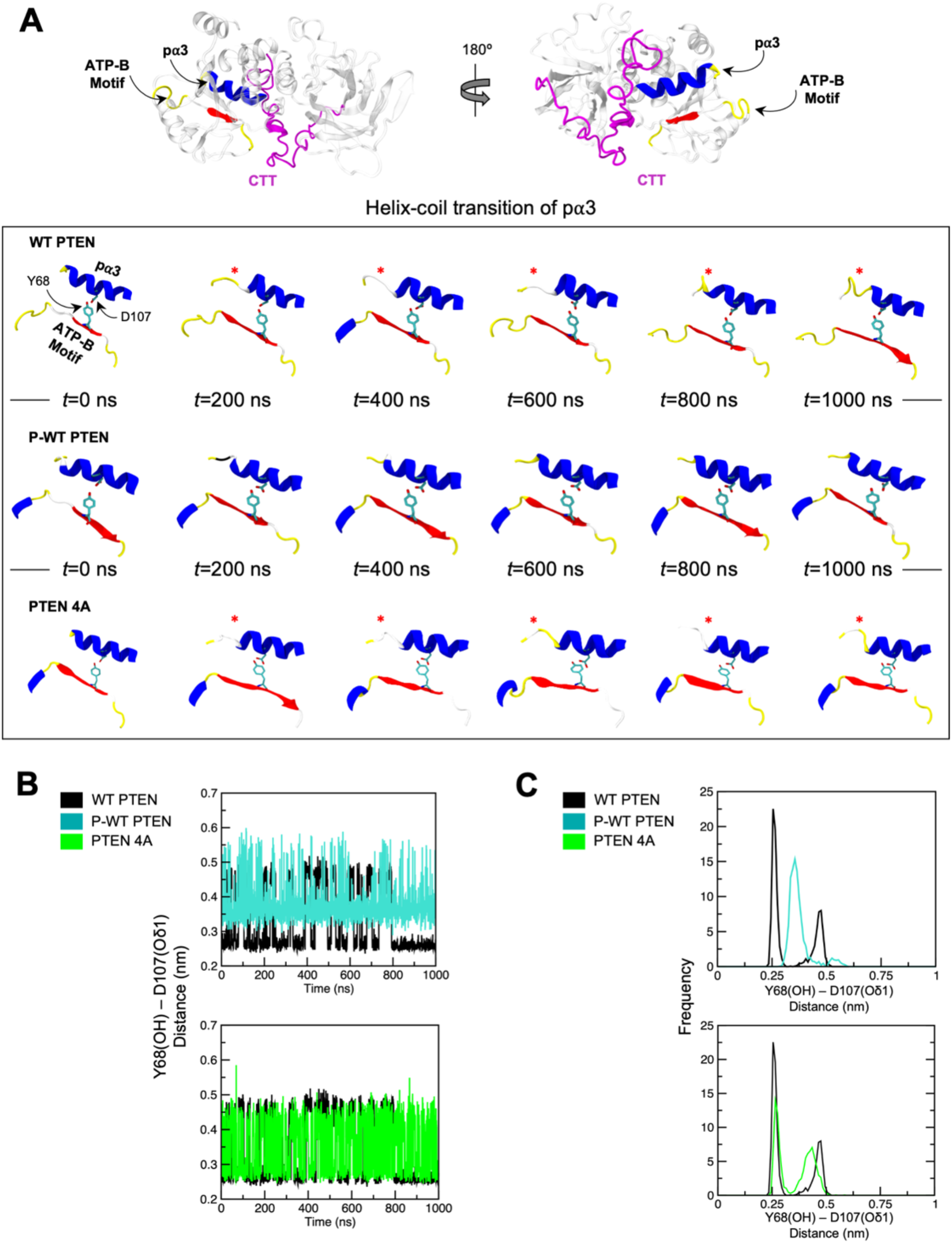
Disruption and rescue of pα3 helix by CTT phosphorylation. Inset highlights the ATP-B binding motif, pα3 helix, and CTT (*magenta*) regions in the PTEN structure. Interacting residues Y68 and D107 are colored in licorice representations. Snapshots taken at 0, 200, 400, 600, 800, and 1000 ns throughout entire simulation for (**A**) WT PTEN (*upper* panel), P-WT PTEN (*middle* panel), and PTEN 4A (*lower* panel). (**B**) Average distance calculated for Y68(OH)-D107(OD1) atoms throughout entire WT PTEN (*black*), P-WT PTEN (*cyan*), and PTEN 4A (*green*) simulations. (**C**) Frequency distribution for WT PTEN (*black*), P-WT PTEN (*cyan*), and PTEN 4A (*green*). Red asterisk (*) indicates location of α-helical unwinding.

**Table 3.** Hydrogen bond occupancy and average distance calculated for Y68(OH)-D107(Oδ1) from MD simulations.

| Simulation Systems | Acceptor | Donor | Frames | Occupancy | <sup>a</sup>D<sub>1</sub> (Å) |
| --- | --- | --- | --- | --- | --- |
| WT PTEN | D107 (OD1) | Y68 (OH) | 2002 | 0.55 | 3.35 (0.9) |
| P-WT PTEN | D107 (OD1) | Y68 (OH) | 2002 | 0.41 | 3.72 (0.5) |
| PTEN 4A | D107 (OD1) | Y68 (OH) | 2002 | 0.42 | 3.55 (0.7) |

<sup>a</sup>D<sub>1</sub> = Avg. distance of Y68(OH)-D107(OD1). Standard deviations are given in parenthesis. Approximately 50,000 frames, each representing a 2 fs time step were utilized for analysis.

### CTT Phosphorylation Induces Structural Changes in the Catalytic Active Site

The position of critical active site residues Asp92 (D92), Cys124 (C124), and Arg130 (R130) is essential for their interaction with PIP_3_ lipid substrate and the overall function of PTEN^87^. Specifically, C124 and R130 are important for the coordination of PIP_3_, whereby R130 facilitates the migration of a phosphate group from the 3’ region of the inositol head group of PIP_3_ to the side chain of C124. Previous studies showed that missense *PTEN* mutations in these active site residues result in severe reduction of phospholipid phosphatase activity of PTEN^2,9,88^. Therefore, active site region, we quantified the interaction distances between D92, C124, and R130 throughout MD simulations (**Figure S10A and S10B**). We found the average distance between the Oδ1 atom of D92 and the Nε atom of R130 (salt-bridge) was 4.9 (0.8) and 4.0 (0.5) Å for WT PTEN and P-WT PTEN, respectively (**Figure S10A**, *left* panel), and P-WT PTEN had a higher frequency of maintaining the salt-bridge than WT PTEN (**Figure S10A**, *right* panel). PTEN 4A behaved similarly to WT PTEN: we observed the average distance between the Oδ1 atom of D92 and the Nε atom of R130 (salt-bridge) to be at 4.6 (0.8) Å, with a slightly higher overall frequency of interaction than WT PTEN (**Figure S10A**, *right* panel). Additionally, a hydrogen bond interaction between C124 and R130 was found to be somewhat similar in all three systems, yet slightly closer in the P-WT PTEN system. The average distance between the Sγ atom of C124 and NH atom of R130 remained unchanged in both WT PTEN and P-WT PTEN systems (**Figure S10B**, *left* panel). However, the C124(Sγ)-R130(NH) hydrogen bond interaction in P-WT PTEN had a slightly higher frequency of interaction than WT PTEN (**Figure S10B**, *right* panel). In contrast, the C124(Sγ)-R130(NH) hydrogen bond in the PTEN 4A system had a slightly lower frequency of interaction than both WT PTEN and P-WT PTEN (**Figure S10B**, *left* and *right* panels). Overall, our simulation results suggest that CTT phosphorylation induces a long-range conformational rearrangement that stabilizes the active site.

Quite often, structural rearrangements of the active site geometry are accompanied or facilitated by conformational changes of secondary structural elements in proximity to the active site. This includes entire a-helices, which often precede or follow a flexible and disordered “lid” region – a characteristic example is the cap domain of a haloalkane dehalogenase^86^. Backbone superimposition of representative centroid structures of P-WT PTEN and PTEN 4A onto WT PTEN reveals phosphorylation-induced CTT conformational changes, which in turn affected the exposure of the active site region (**Figure 5A and 5B**). This long-range conformational rearrangement mediates the reorientation of the catalytic site TI loop which acts as a “lid” to the active site. In the P-WT PTEN structure, the TI loop is rearranged, exposing the WPD and P loops to the solvent (**Figure 5B**), which supports previous studies that the TI loop plays a role in stabilizing interactions between the active site and its PIP_3_ lipid substrate^17,89^. Moreover, the PTEN 4A system also demonstrated an increase in total solvent-accessible surface area (SASA) compared to WT PTEN, yet slightly lower than P-WT PTEN. This suggests CTT mutation at positions S380/T382/T383/S385 to alanine affects TI loop dynamics, further emphasizing the role the CTT has in long-range communication with the active site (**Figure 5C**). Notably, we found that the total SASA substantially increases in the P-WT PTEN system as a result of CTT phosphorylation and may explain the extended conformation of the TI loop (**Figure 5D**). In this context, the TI loop modulates accessibility of the catalytic active site, which suggests CTT phosphorylation may have an impact on the dynamics of the TI loop. Overall, our simulations indicate that upon CTT phosphorylation, a long-range conformational rearrangement mediates the reorientation of the TI loop, exposing the catalytic residues D92, C124, and R130 to the bulk solvent, which is in accordance with the slight displacement of the TI loop (**Figure 5D**).

**FIGURE 5.**
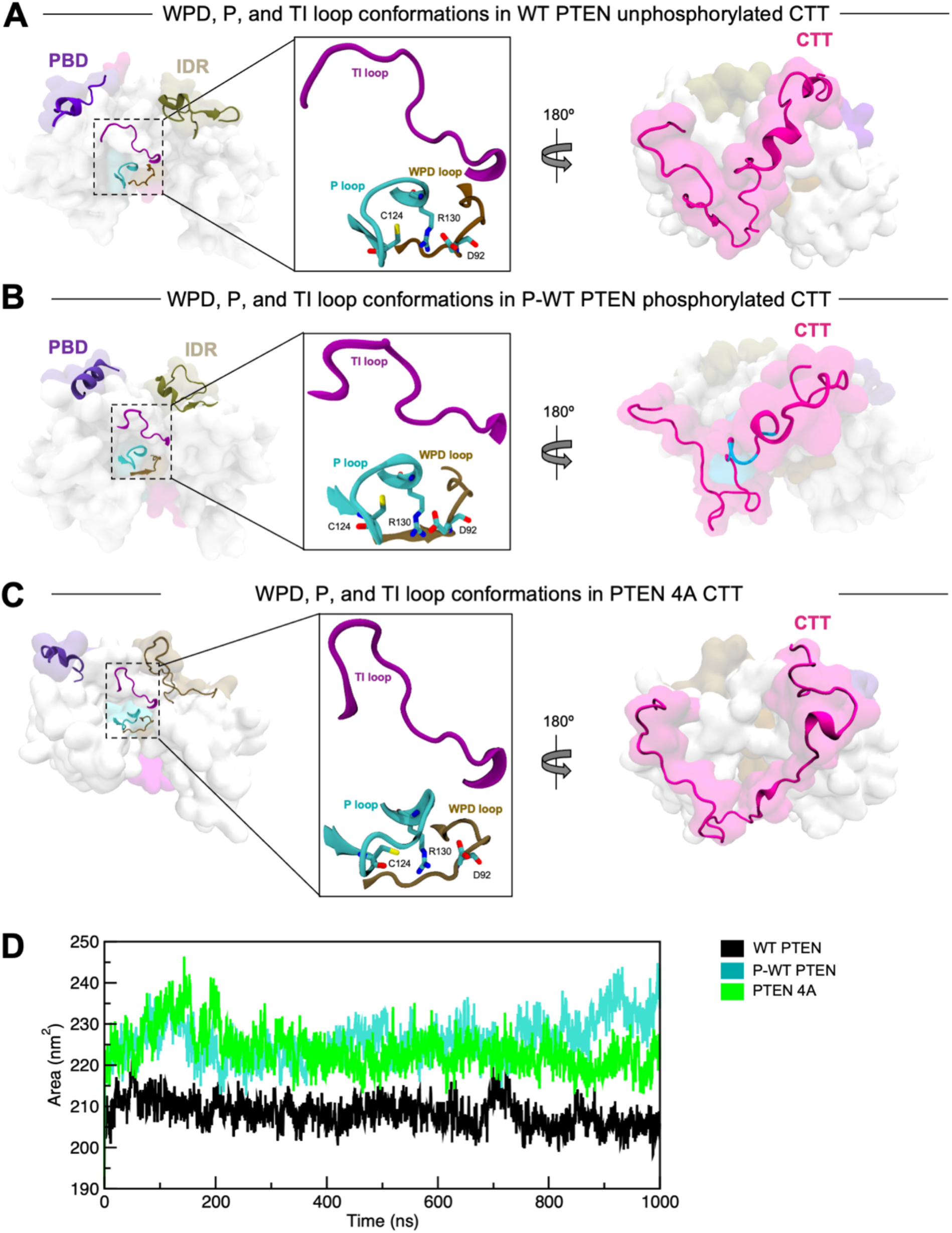
Long-range active site conformational rearrangements. (**A**) WT PTEN, (**B**) P-WT PTEN, and (**C**) PTEN 4A. Inset illustrates D92, C124, and R130 residue positions in sticks within the WPD (*brown*), P (*turquoise*), and TI (*purple*) active site loops. Phosphorylated CTT cluster sites S380/T382/T383/S385 are depicted as surface representations (*light blue*). (**D**) Total solvent accessible surface area (SASA) of WT PTEN (*black*), P-WT PTEN (*cyan*), and PTEN 4A (*green*).

### CTT Phosphorylation Induces Coupling within Interdomain Region and Hydrophobic Region Adjacent to Active Site

The phospho-dependence of PTEN activity^13,18,20–25,30–33^ suggests the existence of allosteric networks across PTEN. Previously, we identified long-range structural communication and a potential basis for allosteric communication formed by the PD-C2D interdomain region of PTEN^90^. Therefore, we sought to identify residues responsible for the long-range transmission of allosteric signals and applied the perturbation response scanning (PRS) method^66^. Anisotropic network models (ANMs)^61,63^ from the representative centroid structures of WT PTEN, P-WT PTEN, and PTEN 4A were constructed using the *ProDy* software package^64^, representing the proteins as elastic mass-and-spring networks. Each PRS map was constructed using the covariance matrices from the first 20 nonzero ANM modes. PRS maps provide information on both the influence of a given (effector) amino acid in transmitting signals when subjected to a perturbation and the sensitivity of a given (sensor) residue to those signals (**Figure 6A, 6C, and 6E**). The effectors clustered in the core and PD-C2D interdomain regions for each system, while the most affected sensors clustered within the disordered PBD, IDR, and CTT regions (**Figure 6B, 6D, and 6F**). For the P-WT PTEN system, a strong coupling extends into residues within the adjacent hydrophobic site (AHS) and ATP-B binding motif of the phosphatase domain, as well as into residues of the Cβ1/2 loop and IDR of the C2 domain, indicating that these regions efficiently propagate structural perturbations to sensor residues induced by CTT phosphorylation (**Figure 6C and 6D**).

**FIGURE 6.**
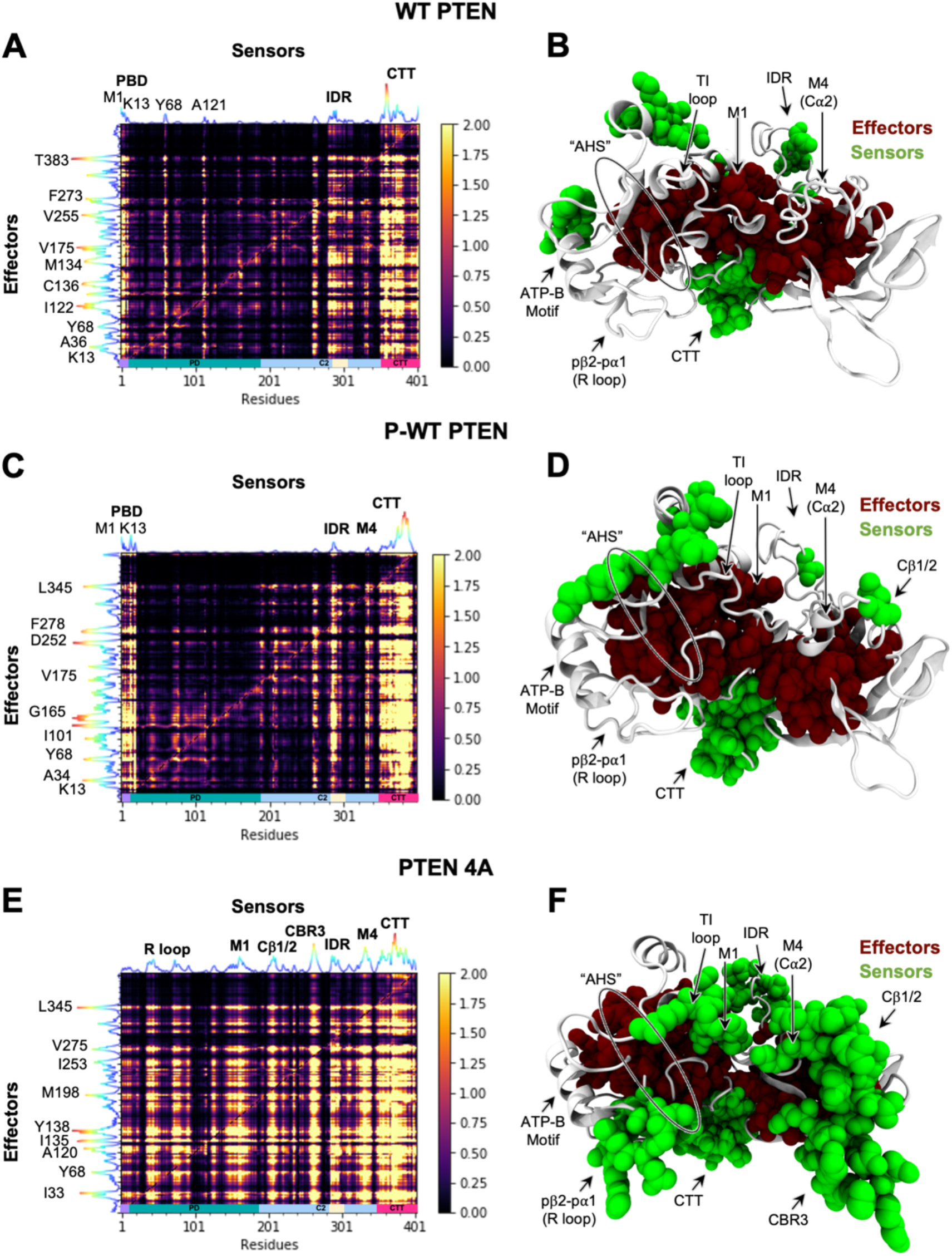
Perturbation response scanning (PRS) analysis identifies highly influential and sensitive residues that likely propagate allosteric signals in WT PTEN, P-WT PTEN, and PTEN 4A. PRS maps (**A, C, and E**) indicate strongest perturbation response sites as shown by the brightest colors (see scale on *right*). The peaks in the curves along the axes indicate the effectors (*left* ordinate) and sensors (*upper* abscissa). Effector (*maroon*) and sensor (*green*) residue sites are indicated by color-coded space filing representation in each of the respective panels for (**B**) WT PTEN, (**E)** for P-WT PTEN, and (**F**) for PTEN 4A. PTEN domains are color-coded along the *lower* abscissa highlighting PBD (*purple*), PD (*cyan*), C2 domain (*light blue*), IDR (*tan*), and CTT (*magenta*) as in **Figure 1**.

Further, for the PTEN 4A system, the PD-C2D interdomain effector residues exerted a remarkably strong influence on sensor residues throughout the entire protein structure, specifically within the pβ2-pα1 loop, TI loop, M1 loop, IDR and three C2 domain loops critical for membrane binding (Cα2, Cβ1/2 and CBR3) (**Figure 6E and 6F**). These results confirm that these regions are highly sensitive to perturbations at CTT residue positions S380/T382/T383/S385 and indicate the bridging role between the CTT and PD-C2 region effectors. In a recent NMR-derived model using VSP (PTEN homolog), Dempsey *et al*.^18^ revealed a phospho-C-tail positioned close to both the CBR3 and Ca2 loops, which confirms our results and supports previous biochemical and structural CTT phospho-dependent interactions with C2 domain^24,25,30,33^. Our results demonstrate the central location of the effectors and their strong influence on sensor residues suggest a role in establishing allosteric communication across the PD-C2D interdomain interface as seen in our previous studies^49,90^.

Next, we identified sites strongly coupled to the active site and CTT that are critical for function and are referred to as dynamic allosteric residue coupling (DARC) sites^91,92^. Interestingly, a mutation at a DARC site is likely to influence conformational dynamics and allosteric regulation^92,93^. To capture the strength of the displacement response of a given residue upon perturbation, we utilized the dynamic coupling index (DCI) metric^93^ within *ProDy* to quantify the long-range coupling within the active site (P loop) and CTT for each system (WT PTEN, P-WT PTEN, and PTEN 4A). We observed that the coupling of the P loop with the AHS, ATP-B motif, TI loop, and pβ2-pα1 (R loop) of the phosphatase domain increased dramatically between WT PTEN and P-WT PTEN (**Figure S11A and 11B**). The CTT demonstrates a similar dynamic coupling pattern with the AHS between WT PTEN and P-WT PTEN, though not as strongly coupled as the P loop (**Figure S11A and 11B**). Previous studies have also shown that the AHS further stabilizes the conformation of PTEN needed for optimal processive catalysis^37^. Thus, the increase in the dynamic coupling of the active site (P loop) with the AHS in the P-WT PTEN system may be linked with lower catalytic activity and represents a critical allosteric mechanism of PTEN. For the PTEN 4A system we found coupling of the P loop to be enhanced with the residues in the AHS, IDR, and core of the C2 domain, whereas a dramatic increase in coupling of the CTT is seen throughout the entire protein structure (**Figure S11C**), consistent with the regions identified in PRS analyses (**Figure 6E and 6F**). Previous reports indicate that PTEN 4A overrides autoinhibitory activity^21,94^, which increases membrane localization^33,81,82^ to produce a pronounced tumor suppressive effect due to its enhanced lipid phosphatase activity^95^. Importantly, in a recent study PTEN 4A acts as a gain of function variant causing a substantial increase of WT PTEN function^94^. When comparing the DCI profiles of the catalytic P loop and CTT across each system, fluctuations in DCI imply that the long-distance communication pathways do not follow similar channels to the active site and may be the result of altered dynamics (**Figure S11D**). These results suggest that positions exhibiting higher dynamic coupling to the active site and CTT may have a greater impact on protein function. Our data also suggests a link between positions with high dynamic coupling and regions critical for allosteric regulation. These results support our previous *in silico* computational data^49,90^ and experimental findings, which originally identified the AHS as another region that can play a role in monomer ligand binding^37,89,96^.

To shed further light on this hypothesis, potential allosteric binding pocket sites were detected using *ProDy* essential site scanning analysis (ESSA)^67^, an elastic network model-based method that detects allosteric sites that modulate dispersion of protein global motions. ESSA incorporates *Fpocket*^68^ for the detection of pockets that potentially bind small ligands. As shown in **Figure 7A and 7B**, detection of potential binding pockets in the PD-C2D interdomain region in both WT PTEN (pockets A1, A4, and A5) and P-WT (pockets B1 and B5), as well as the AHS region in both WT (pocket A2) and P-WT (pocket B4) PTEN systems, indicate that these regions may form essential allosteric sites, which may be used for the design of potential PTEN inhibitors. As indicated in **Table 4**, residues within the AHS are underlined for WT PTEN (pocket A2), P-WT PTEN (pocket B4), and PTEN 4A (pocket C2) systems and common pocket residues within the pockets of each system are indicated in bold. The pockets are numbered by rank according to their allosteric potential. The detection of the top five pockets and potential allosteric residues is similar across the WT PTEN and P-WT PTEN systems - specifically, pocket 2 (A2) in WT PTEN matches pocket 4 (B4) in P-WT PTEN; pocket 4 (A4) in WT PTEN matches pocket 1 (B1) in P-WT PTEN; pockets 5 (A5/B5) are the same in both systems; and pocket 3 (A3) in WT PTEN matches pocket 2 (B2) in P-WT PTEN. However, the volume of pocket A3 is slightly decreased from 673.17 Å^2^ in WT PTEN to 663.60 Å^2^ in pocket B2 of P-WT PTEN, indicating that CTT phosphorylation slightly diminishes the allosteric binding capacity in this region (**Table 4**). Notably, the fifth most important pocket 5, which lies within the PD-C2D interdomain region is substantially diminished in volume in P-WT PTEN (pocket B5) compared to WT PTEN (pocket A5), 472.15 and 745 Å, respectively (**Table 4**). Of interest is the detection of pocket 3 (B3) proximal to the CTT in P-WT PTEN, indicating that there is an increase in essentiality of residues in this region, and/or an enhanced binding capacity, either of which being likely to have allosteric implications. Additionally, pocket 1 of the WT PTEN system (A1) is lost in the P-WT PTEN system, suggesting that the CTT phosphorylation cluster may disrupt this allosteric binding site in P-WT PTEN.

**FIGURE 7.**
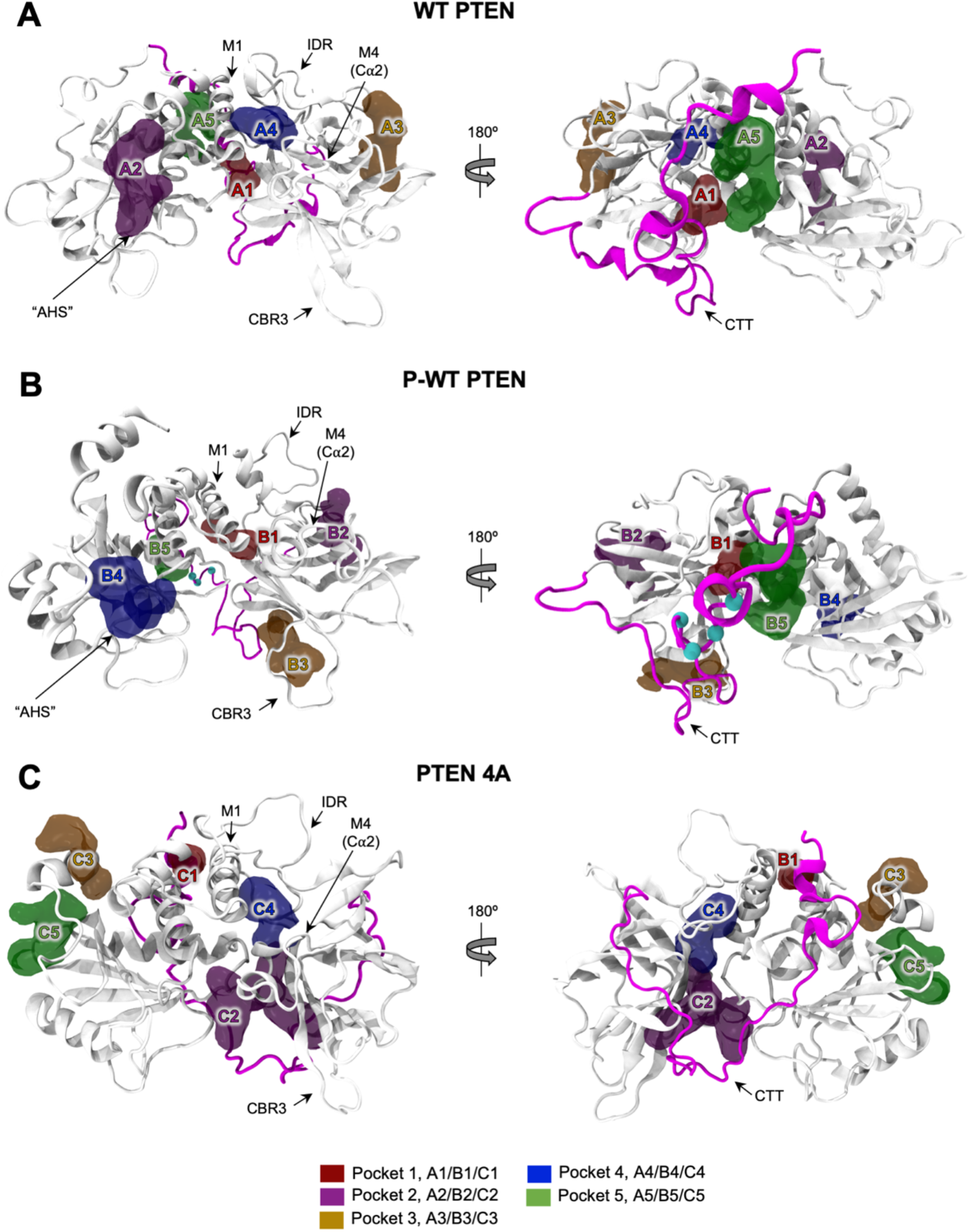
Top five potential allosteric pockets for (**A**) WT PTEN, (**B**) P-WT PTEN, and (**C**) PTEN 4A. Pockets are ranked based on their allosteric potential: Pocket 1, A1/B1/C1 (*red*), Pocket 2, A2/B2/C2 (*purple*), Pocket 3, A3/B3/C3 (*orange*), Pocket 4, A4/B4/C4 (*blue*), and Pocket 5, A5/B5/C5 (*green*). The CTT is highlighted in *magenta* in the PTEN structures. Phosphorylated CTT cluster sites S380/T382/T383/S385 are depicted as spheres (*light blue*).

**Table 4.** Residues in top five allosteric pockets. Pockets 1, 2, 3, 4, and 5 are indicated in *red, purple, orange, blue*, and *green*, respectively, in **Figure 7**.

| System | Pocket 1<br>(A1/B1/C1) | Pocket 2<br>(A2/B2/C2) | Pocket 3<br>(A3/B3/C3) | Pocket 4<br>(A4/B4/C4) | Pocket 5<br>(A5/B5/C5) |
| --- | --- | --- | --- | --- | --- |
| WT PTEN | N94, **P95**, Q97, **L98**, E99, **Y188**, Q219, L220, C250, G251, **D252**, **W274**, **N276**, R378, Y379, S380 | **D24**, L25, T26, V45, **R47**, N48, <u>**A126**</u>, G127, <u>**K128**</u>, Y155, R159, T160, G165 | **K197**, **M198**, **M199**, T232, **R233**, F238, **Y240**, **F241**, **E242**, **E307**, **A309**, **D310**, **Y315**, **Y346** | P169, R172, **R173**, **Y177**, **W274**, **V275**, **F279**, L295, I300, **I303**, C304, **L318**, L320, N323, **D324** | L98, **E99**, **K102**, **P103**, **E106**, **L140**, H141, R142, Y178, L181, **L182**, L186, D187, Y188, **D395**, Q396, **Q399**, **I400** |
| Total Volume | 276.84 Å<sup>2</sup> | 605.76 Å<sup>2</sup> | 673.17 Å<sup>2</sup> | 392.60 Å<sup>2</sup> | 745.00 Å<sup>2</sup> |
| P-WT PTEN | **P95**, **L98**, **S170**, **R173**, **Y174**, **Y177**, Y178, L181, **Y188**, **D252**, **W274**, V275, **N276**, F279 | **K197**, **M198**, **M199**, **R233**, **Y240**, **E242**, **E307**, R308, **A309**, **D310**, N311, E314, **Y315**, **F241**, K344, **Y346** | Q214, V216, **K223**, I224, Y225, S226, **E256**, F258, H259, Q261, M264, L265, K266, K267, D268, K269, S368, **V369**, D371 | Q17, E18, D19, L23, **D24**, G36, F37, V45, <u>**R47**</u>, K125, <u>**A126**</u>, <u>**K128**</u> | **E99**, **K102**, **P103**, **E106**, pT383, pS385, D386, N389, **D395**, **Q399**, |
| Total Volume | 293.40 Å<sup>2</sup> | 663.60 Å<sup>2</sup> | 795.37 Å<sup>2</sup> | 474.60 Å<sup>2</sup> | 472.15 Å<sup>2</sup> |
| PTEN 4A | **L140**, L146, S179, **L182**, H397, **I400**, K402 | A72, E73, R74, E91, D92, H93, N94, <u>**K128**</u>, V216, C218, L220, **K223**, D252, K254, V255, **E256**, H272, F273, W274, L320, D324, D326, N329, K330, **V369**, S370, N372, E373, P374, D375, R378 | I5, K6, E7, I8, V9, S10, R11, R14, Y16, I28, Y29, F154, E157, V158 | H93, P95, **S170**, **R173**, **Y174**, **Y177**, **W274**, **V275**, **F279**, I280, G282, P283, **I303**, S305, **L318**, T319, **L320**, **D324** | V9, N12, R15, Y27, P30, K60, H61, D116, N117, V119 |
| Total Volume | 167.42 Å<sup>2</sup> | 1256.09 Å<sup>2</sup> | 466.32 Å<sup>2</sup> | 506.95 Å<sup>2</sup> | 597.54 Å<sup>2</sup> |

Pockets are numbered by rank according to their allosteric potential. Residues within the AHS are underlined for WT PTEN (pocket A2), P-WT PTEN (pocket B4), and PTEN 4A (pocket C2) systems. Common pocket residues within each system are indicated in bold.

Next, we determined if CTT mutation at positions S380/T382/T383/S385 to alanine (PTEN 4A) affects the detection of possible allosteric binding pockets compared to WT PTEN and P-WT PTEN systems. The detection of two PD-C2D interdomain pockets are similar across the WT PTEN and PTEN 4A systems - specifically, pocket 4 (A4) in WT PTEN matches pocket 4 (C4) in PTEN 4A and pocket 5 (A5) in WT PTEN matches pocket 1 (C1) in PTEN 4A. However, the volume of pocket A4 is markedly increased from 392.60 Å^2^ in WT PTEN to 506.95 Å^2^ in pocket C4 of PTEN A4, indicating enhanced allosteric binding capacity in this region (**Table 4**). Of significance are two distinct pockets detected in PTEN 4A (pockets C3 and C5), suggesting that the phosphorylation-deficient effect of the CTT is likely coupled to the enhanced allosteric binding capacity in this region (**Figure 7C**). These results are also consistent with the regions identified in both the perturbation response scanning and dynamics coupling analyses (**Figure 6E and 6F; Figure S11C**). Our allosteric pocket analyses identified residues in the AHS consistent with previous reports, which describe this site as a putative binding site apart from the active site (**Figure 7A and B; Table 4**)^37,89,96^. Notably, the sites identified to have the strongest perturbation response and thus critical for long-range structural communication of CTT are consistent with findings from other research groups. Interestingly, these regions are also involved in membrane binding, as identified by us and others^33,39,97^, suggesting that these sites are coupled to the CTT. Indeed, a direct interaction between the CTT and the PBD was revealed by hydrogen/deuterium exchange (HDX) mass spectrometry (MS) site directed mutagenesis and photo cross-linking experiments where the phosphorylated CTT interacts with both the catalytic and C2 domains^30^.

### Exploration of PTEN CTT-PDZ domain interactome and PTEN CTT-PDZ complexes

PTEN dynamic regulation and stability is coupled to its CTT phosphorylation state^13,18,20–25,30–33^. Aside from CTT phosphorylation, the two putative PEST sequences (residues 350-375 and 379-386) and PDZ binding motif (401TKV403), mediate crucial PPIs and further regulate PI3K/AKT/mTOR cell signaling pathways^13,14,20,21,32,98,99^. PDZ domains are a class of modular interaction domains, generally part of larger proteins, that bring binding partners together into complexes^100^. PTEN has multiple binding partners containing PDZ domains^101^; some of which are important in cell-specific ways, such as DLG4 in the synapse. In **Figure 8A**, we have illustrated known PDZ-domain-containing interactors with the CTT of PTEN including both Glycogen Synthase Kinase 3-beta (GSK3β) and Casein Kinase 2 (CK2), kinases that are known to phosphorylate the CTT^21,102–105^. Moreover, we performed a network analysis on these known CTT interactors, including on the known relationships between them as catalogued by the STRING database^73^. We found that by betweenness centrality measures DLG4 and MAGI3 are possibly the most vital to coordinating PTEN protein-protein interactions (**Figure 8A**). The expansion in node size and progression toward red color are proportional to its degree and importance (betweenness centrality) in the network, respectively. Beyond DLG4 and MAGI3, there are many other interesting biological PTEN CTT interactors that have been subject to little study, especially in terms of affecting PTEN structure-function dynamics^13,14^. For instance, MAST2’s PDZ domain binds to PTEN, stabilizing the protein and phosphorylating PTEN at an unknown site^101^. In contrast, PTEN binding to MAGI2’s second PDZ domain is reduced when PTEN is phosphorylated at its CK2 phosphorylation sites within the CTT cluster (residues 380-385)^21,102^.

**FIGURE 8.**
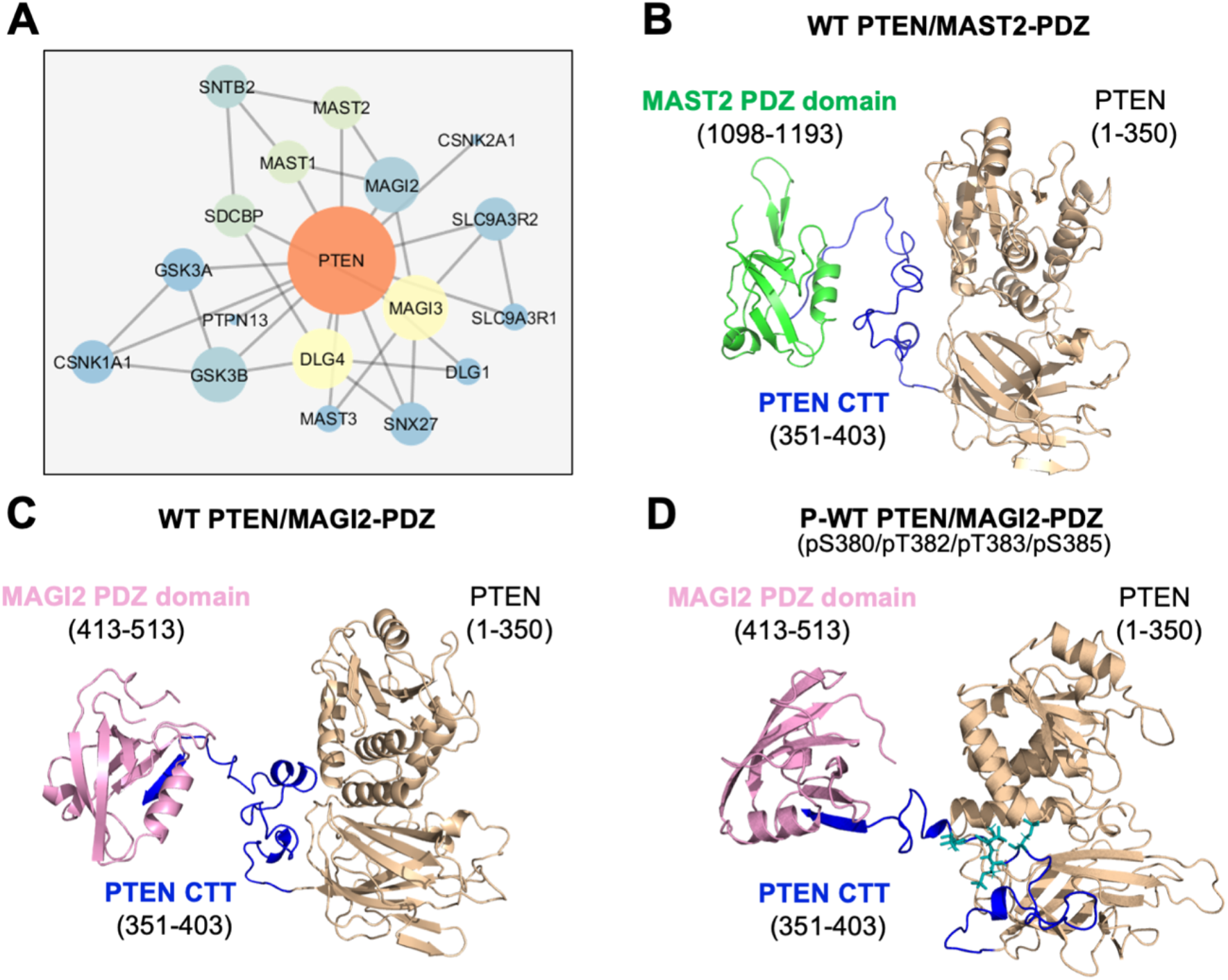
PTEN-PDZ PPI network and *in silico* comparison between docked complexes. (**A**) Putative PTEN-PDZ PPI network, where each node represents a protein, and each edge (line) refers to an interaction. The expansion in node size and progression toward red color are proportional to its degree and importance (betweenness centrality) in the network, respectively. Docked complexes of (**B**) WT PTEN/MAST2-PDZ, (**C**) WT PTEN/MAGI2-PDZ, (**D**) P-WT PTEN/MAGI2-PDZ. MAST2-PDZ domain (*green*), MAGI2-PDZ domain (*pink*), PTEN CTT (*blue*), and PTEN structure (*tan*) are color-coded accordingly.

To demonstrate PTEN PPIs involving the CTT and PDZ domains, we constructed models for WT PTEN/MAST2 PDZ, WT PTEN/MAGI2, and P-WT PTEN/MAGI2 complexes using the full-length PTEN models and available NMR structures of PDZ domains (MAST2 PDZ: PDBID 2KYL; MAGI2 second PDZ: PDBID 1UEQ; see Methods) [**Figure 8B, 8C, and 8D**]. In **Figure 8D**, the P-WT PTEN/MAGI2-PDZ model reveals a “partially collapsed” conformation against the PD-C2 interdomain interface in P-WT PTEN, which is consistent with our MD simulations (**Figure 2D**; **Figure 5B; and Figure S6**). In general, PDZ domains bind to PDZ-binding motifs that contain the C-terminal residues of their binding partners. The motifs are incorporated as anti-parallel β-strands between the helix α_B_ and strand β_B_ of the domain with binding specificity between partners determined by additional interactions^100^. The PDZ-binding motif was not in the freely available, β-strand conformation needed to bind a PDZ domain in the full-length PTEN models. To overcome this difficulty, residues 1-390 of PTEN were fused using PyMOL^55^ to the β-strand-like PTEN motif peptide from the WT PTEN motif/MAST2 PDZ complex (PDBID: 2KYL, PTEN residues 391-403) [**Figure 8B**]. Models were constructed and refined using HADDOCK and Rosetta3^77,78^ and their quality evaluated with MolProbity^79^ (**Table 1**). PTEN PDZ-binding motif was incorporated as an extended polypeptide or β-strand into the MAST2 and MAGI2 PDZ domains, burying between 1025-1218 Å^2^ of surface area in the binding interface (**Figure 8B, 8C, and 8D; Table 1**).

The presence of the four phospho-residues does not prevent *in silico* association between the PTEN PDZ motif and MAGI2-PDZ domain, which was surprising given the phospho-dependence of binding in cellular and biochemical assays^21,102^. It is likely that the solvent accessible surface area (SASA) and structural conformation of the PDZ motif may play a role in the PDZ binding interactions. Indeed, in the WT PTEN and P-WT PTEN MD simulations, the estimated total SASA in the PDZ binding motif was substantially decreased in the P-WT PTEN system (**Figure S12A and Table 5**), implying it is less accessible for binding. The initial PTEN structures used in modeling had the motif already in β-strand conformation. In full-length apo models, WT and P-WT PTEN PDZ motifs are in coil and turn conformations (**Figure 3**), not the β-strand found during PDZ binding. During our MD simulations, PDZ motif residues T401 and K402 of WT PTEN have backbone φ,ψ torsion angles consistent with a β-strand conformation (**Figure S12B**). Conversely, in P-WT PTEN, these residues are more a-helical, which indicate that a phosphorylated state possibly stabilizes a more a-helical state in terms of φ,ψ torsion angles and suggests a possible mechanism for the phospho-dependence of PDZ binding observed in prior experimental studies. Further, during our MD simulations we investigated the dynamics of T401 and K402 side chains in terms of time-evolution of torsion angle χ_1_ (**Figure S12C).** In P-WT PTEN, abrupt fluctuations of the χ_1_torsion angle were observed in T401 within the range of −200° to 200° before finally stabilizing at 50° after 400 ns (**Figure S12C, *left* panel**). However, CTT phosphorylation specifically stabilizes χ_1_ torsion angle of K402 at −50° throughout the entire simulation (**Figure S12C, *right* panel**). Overall, we conclude that CTT phosphorylation may facilitate binding by stabilizing PDZ conformation – though large-scale simulations are necessary to further assess the PTEN CTT-PDZ and MAGI2/MAST2 PDZ binding interactions which is outside the scope of this current study.

**Table 5.** Average SASA of each residue within PDZ-binding motif. Standard deviations are given in parenthesis.

| Simulation Systems | T401 Avg. SASA (Å<sup>2</sup>) | K402 Avg. SASA (Å<sup>2</sup>) | T403 Avg. SASA (Å<sup>2</sup>) |
| --- | --- | --- | --- |
| WT PTEN | 7.11 (1.6) | 17.63 (3.5) | 12.83 (4.8) |
| P-WT PTEN | 0.98 (0.9) | 9.55 (2.5) | 11.88 (3.3) |

## CONCLUSION

Overall, our current work employs an *in silico* theoretical framework based on empirical data. Our results reveal the mechanistic interplay of CTT phosphorylation, conformational dynamics, and coupled underpinning of allosteric regulation in PTEN, demonstrating how changes to the S380/T382/T383/S385 phosphorylation sites in the CTT can cause long-range changes to PTEN catalytic and allosteric AHS sites. Additionally, we illustrate how the phosphorylation state may affect the ability of PTEN PDZ motif to bind to PTEN’s binding partners. Although our studies provide a structural basis for further understanding the mechanism of PTEN CTT phosphorylation and offers further insight on the regulation of PTEN, additional work is needed to discern the effects of different germline *PTEN* mutations on CTT phosphorylation. Further, the CTT enzymatic regulation and localization represent modalities that may be tuned differentially for explaining the clinical heterogeneity associated with different *PTEN* mutations. The current work provides insight into how PTEN transmits signals from CTT to other regions in the structure, and it provides a framework to connect extensive experimental studies needed to support our present work. Altogether, our study provides mechanistic understanding of the detailed atomistic interactions and structural consequences of PTEN CTT phosphorylation and broadens our insights into potential allosteric druggable target sites as a viable and currently unexplored treatment approach for individuals with PHTS.

## Supporting information

Supplemental Script

## ASSOCIATED CONTENT

### Supporting information

Supporting methods, supplemental figures, and tables are available in a separate document, as well as the scripts and commands used for Rosetta FloppyTail model generation.

- Conformational ensemble structures from top two representative clusters of PTEN CTT, Rosetta FloppyTail scores/energies and RMSD for PTEN CTT, Conformational ensemble structures from top two representative clusters of I-TASSER and Rosetta FloppyTail MD simulation replicas, Conformational dynamics of PBD and IDR regions, Conformational dynamics of CTT region, Conformational ensemble structures from MD simulation replicas of IDR and CTT regions, Calculation of a-helical probability, Secondary structure analysis of ATP-B binding motif and pα3, Distance interaction between catalytic residues D92-R130 and C124-R130, Comparison of the coupling of the P loop and CTT regions, Average SASA per residue and comparative analysis of χ_1_ sidechain torsion angle, and Ramachandran plots for PDZ binding motif; and Tables S1-S4: Comparison of the full-length *in silico* PTEN structure (I-TASSER) simulation systems, Reproducibility of the full-length *in silico* PTEN structure (I-TASSER) simulation systems, Sampling replicas from different initial full-length *in silico* PTEN structures, and Sampling replicas from different full-length *in silico* PTEN models (**PDF**)
- PTEN CTT Network Statistics (Table S5) (**XLXS**)
- Scripts and commands used for Rosetta FloppyTail model generation (**DOCX**)

### Data and Software Availability

The GROMACS software necessary for running the MD simulations is publicly available at https://manual.gromacs.org/2020.2/. The *ProDy* software utilized for carrying our ENM, PRS, DCI, and ESSA analyses is publicly available at http://prody.csb.pitt.edu. Usage instructions and documentation for both GROMACS and *ProDy* are provided at respective weblinks. The complete MD trajectories used (a) to plot the *R_g_* time series, (b) to plot the RMSF time series, (c) to plot the RMSD time series, (d) to plot the DSSP distributions, (e) to plot distance distributions, (f) to plot SASA time series, (g) to plot dihedral angle analysis, (h) to plot α-helical propensities, (i) to map DCI profiles, (j) to generate PRS maps, and (k) to generate ESSA pocket analyses are available from the author upon request. The Rosetta FloppyTail script is provided as Supporting Information.

## ACKNOWLEDGEMENT

We thank Ann Tushar, Hyunbum Jang, and Ruth Nussinov for their critical review of the manuscript. This study was funded, in part, by the Ambrose Monell Foundation PTEN-Switch Grant, a grant allocation of computing time from the Ohio Supercomputing Center (PCCF0020) [both to C.E.], the National Institutes of Health (NIH) [P41GM103712 to I.B.] and the Molecular Sciences Software Institute (MolSSI) [COVID-19 Seed Software Fellowship to J.K.]. I.N.S. is funded, in part, by the Ambrose Monell Cancer Genomic Medicine Fellowship, the NIH National Cancer Institute T32 5T32CA59366-22, and NIH National Institute of General Medical Sciences (NIGMS) Maximizing Opportunities for Scientific and Academic Independent Careers (MOSAIC) K99/R00 Grant – 1K99GM143552-01. I.V. is the John K. Vries Chair at the University of Pittsburgh. C.E. is an American Cancer Society Clinical Research Professor and the Sondra J. and Stephen R. Hardis Endowed Chair of Cancer Genomic Medicine at the Cleveland Clinic.

## AUTHOR INFORMATION

### Author Contributions

C.E. and I.N.S. conceived and designed the study. I.N.S. set up, performed, analyzed, and produced visualizations for simulations. J.D. performed, analyzed, and produced visualizations for molecular docking. S.T. performed and analyzed PPI network. I.N.S., J.K., J.D., S.T., I.B. and C.E. interpreted all results. I.N.S., J.D., S.T., and C.E. wrote the manuscript. All authors critically revised the manuscript and gave final approval.

### Notes

The authors declare no competing financial interests.

