## Supporting Information for "Structural and Dynamic Effects of PTEN C-terminal Tail Phosphorylation"

*for*

**C-terminal Tail Phosphorylation**

Iris N. Smith^1^, Jennifer E. Dawson^1^, James Krieger^6^, Stetson Thacker^1,2^, Ivet Bahar^6^, and Charis Eng^1-5*^

^1^Genomic Medicine Institute, Lerner Research Institute, Cleveland Clinic, 9500 Euclid Avenue, NE-50, Cleveland, OH, 44195, USA; ^2^Cleveland Clinic Lerner College of Medicine, Case Western Reserve University, 9500 Euclid Avenue, Cleveland, OH, 44195, USA; ^3^Case Comprehensive Cancer Center, Case Western Reserve University School of Medicine, 10900 Euclid Avenue, Cleveland, OH, 44106, USA; ^4^Taussig Cancer Institute, Cleveland Clinic, 9500 Euclid Avenue, Cleveland, OH, 44195, USA; ^5^Department of Genetics and Genome Sciences, Case Western Reserve University School of Medicine, 10900 Euclid Avenue, Cleveland, OH, 44106, USA ; ^6^Department of Computational and Systems Biology, University of Pittsburgh, 800 Murdoch Building, 3420 Forbes Avenue, Pittsburgh, PA 15260, USA

*Corresponding author

**Supporting Methods**

**Generation of full-length *in silico* PTEN model for additional testing on the effect of the WT PTEN CTT conformation**. The full-length model of PTEN was generated using Rosetta FloppyTail application^1^. Initial structural poses for the disordered CTT of WT PTEN were relaxed with Rosetta3^2-5^ and Rosetta3’s cleanup utility to pre-process the structures. The FloppyTail protocol used the PTEN amino acid sequence, the processed initial structure, and the 3-residue fragment set for PTEN (generated by https://robetta.bakerlab.org/fragmentsubmit.jsp). The CTT PTEN (residues 351-403) were treated as a flexible chain during the FloppyTail application. Ten structures were generated. Their Rosetta scores/energies and RMSD for their CTT domains relative to the initial relaxed structure can be seen in **Figure S3**. The FloppyTail model of PTEN used as an initial structure during molecular dynamics (MD) simulations was further relaxed and equilibrated with the MD software (see **Materials and Methods**) prior to production. The software scripts and commands used to model the CTT are provided as **a separate DOCX file**. The lowest energy model from Rosetta FloppyTail was used as the initial structure for MD simulations (**Figure S4**). All-atom MD simulations were conducted as described in Materials and Methods.

**Supporting Figures**

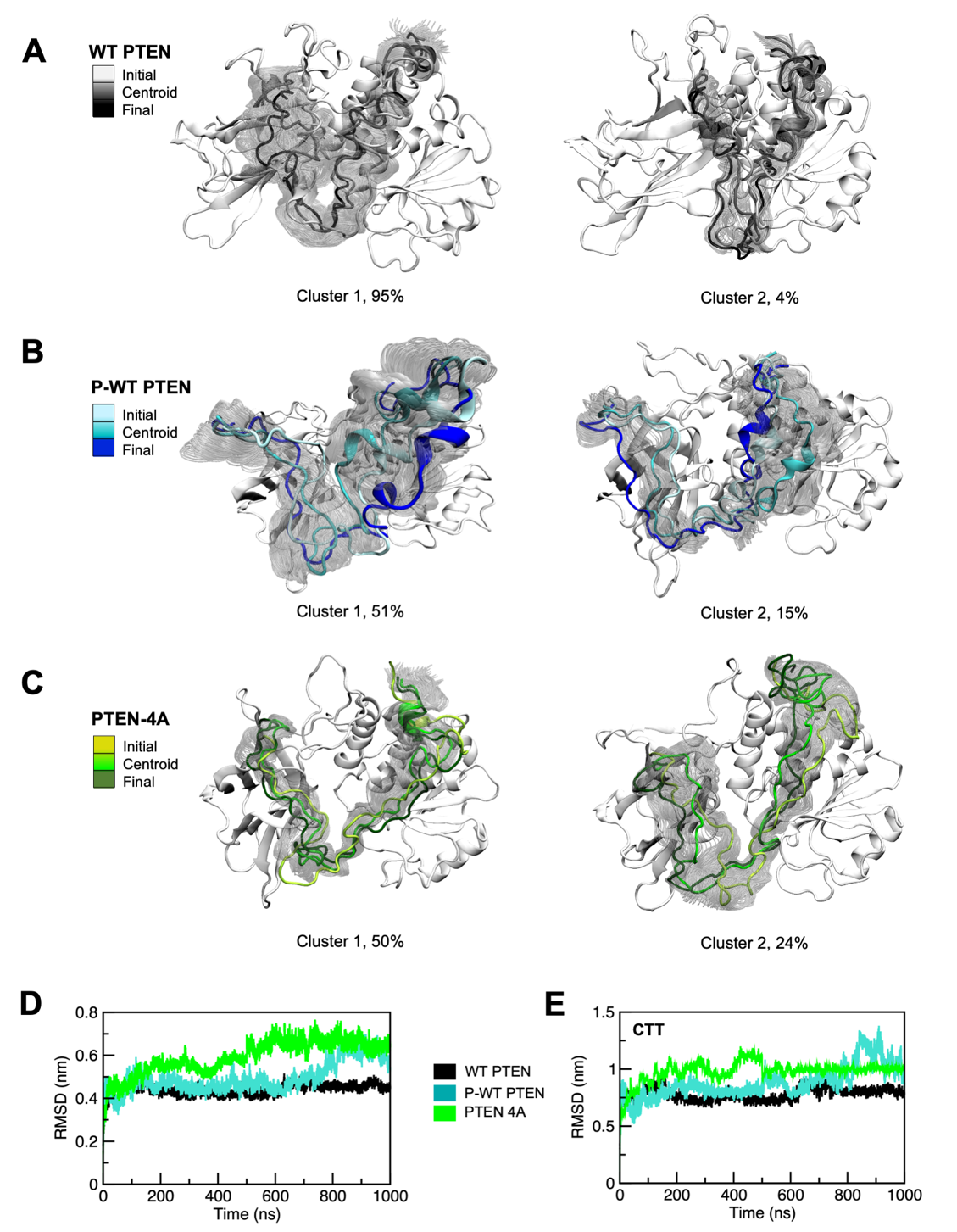

**Figure S1**. Conformational ensemble structures from top two representative centroid clusters of PTEN CTT for (**A**) WT PTEN, (**B**) P-WT PTEN, and (**C**) PTEN 4A. The backbone RMSD time profiles are shown from the initial structure for each system (**D**) and CTT only RMSD for each system (**E**). Initial, centroid, and final conformations are colored as a gradient in *black*, *blue*, and *green* for WT PTEN, P-WT PTEN, and PTEN 4A, respectively. All other structures in each ensemble are indicated in translucent *gray*. Clusters are from the top two dominant conformational ensembles of WT PTEN, P-WT PTEN, and PTEN 4A, with populations at 95%, 51%, and 50%, respectively for cluster 1, and at 4%, 15%, and 24%, respectively for cluster 2.

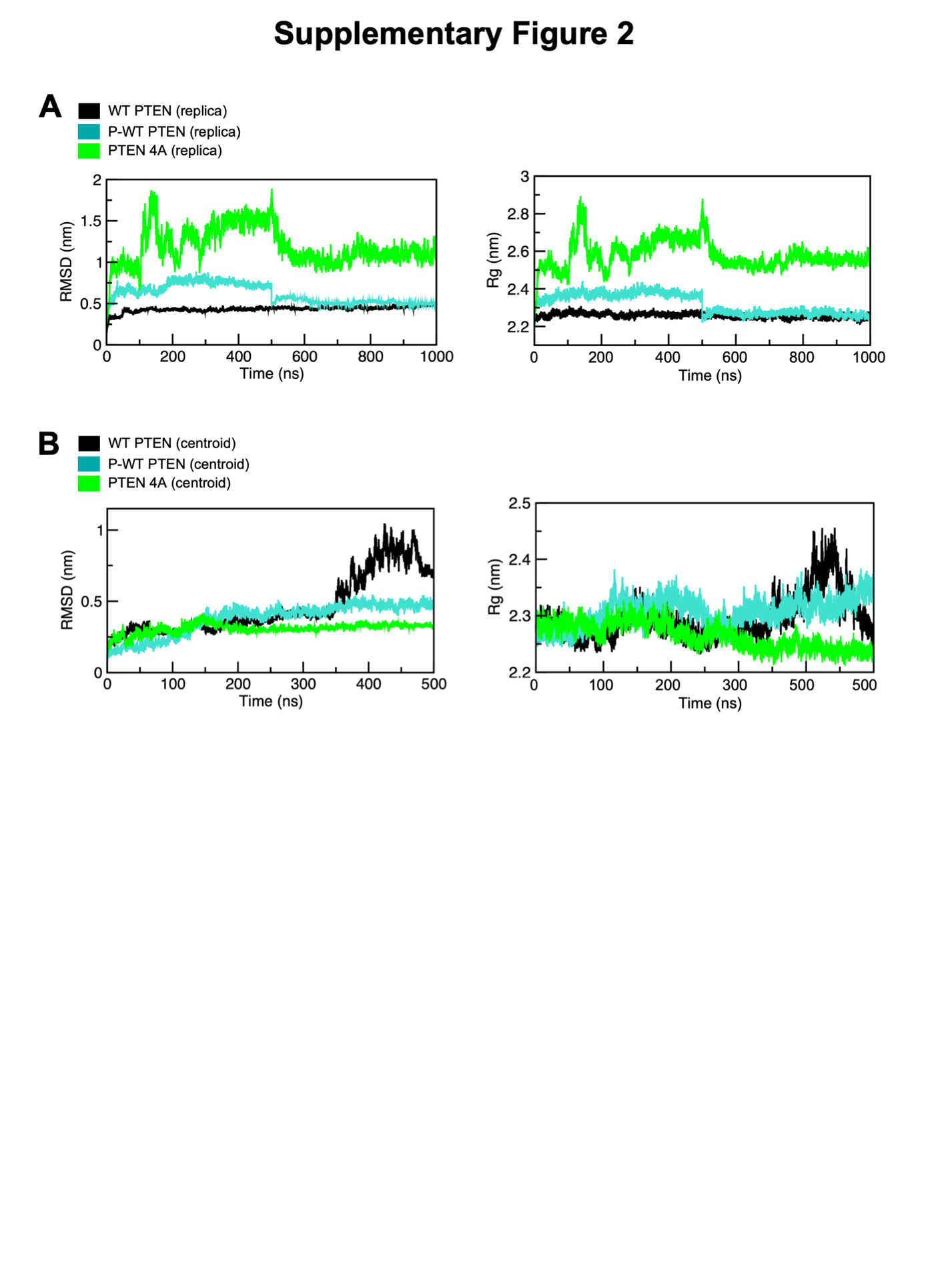

**Figure S2**. Production run reproducibility RMSD time profiles for WT PTEN, P-WT PTEN, and PTEN 4A. The backbone RMSD time profiles for each system, with the 1 μs replica (**A**) and 500 ns centroid structure replicas (**B**) for each system. Each system is colored in *black*, *blue*, and *green* for WT PTEN, P-WT PTEN, and PTEN 4A, respectively.

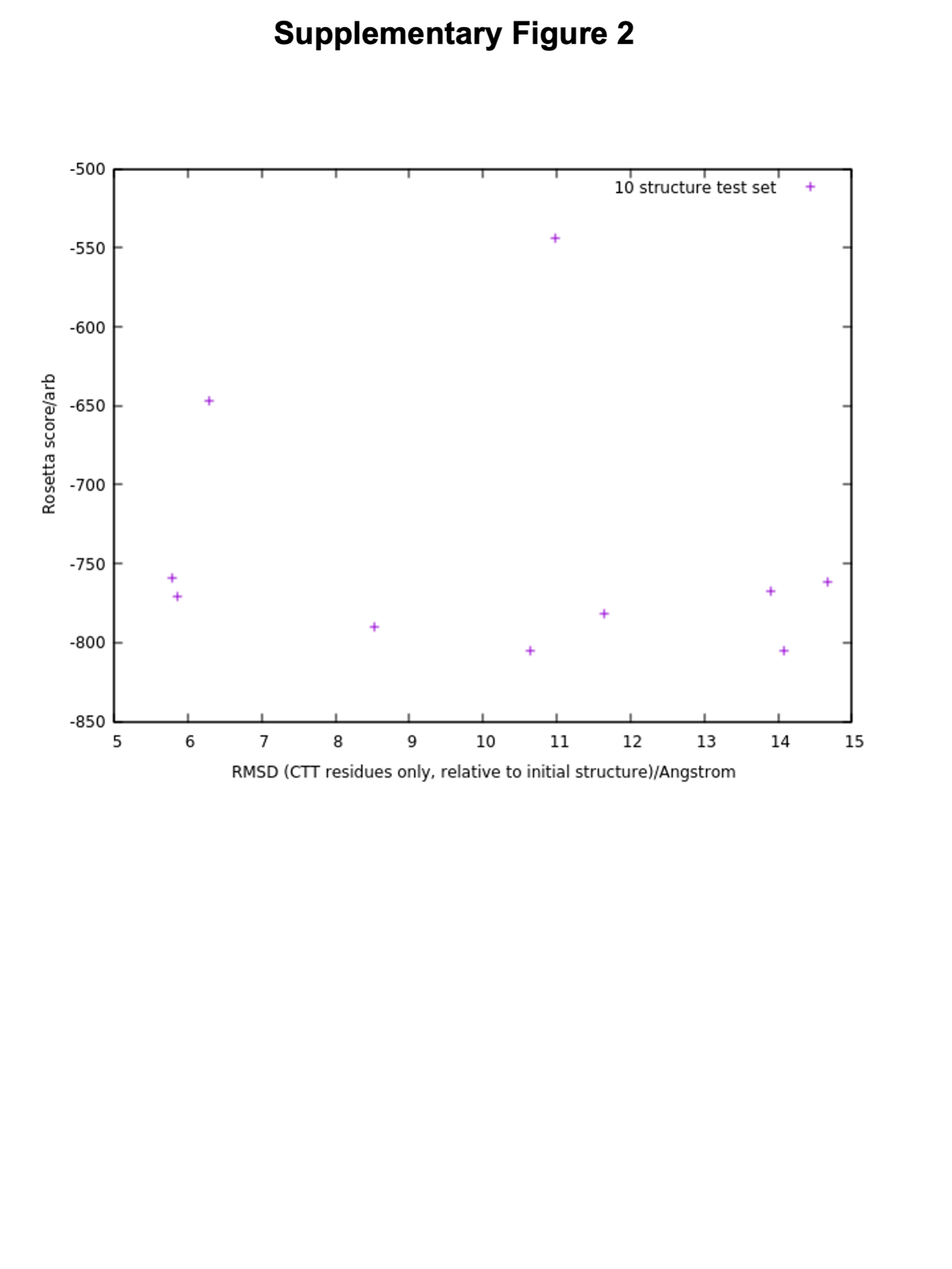

**Figure S3**. Rosetta FloppyTail scores/energies and RMSD for PTEN CTT. Rosetta scores/energies and RMSD for their CTT domains relative to the initial relaxed structure.

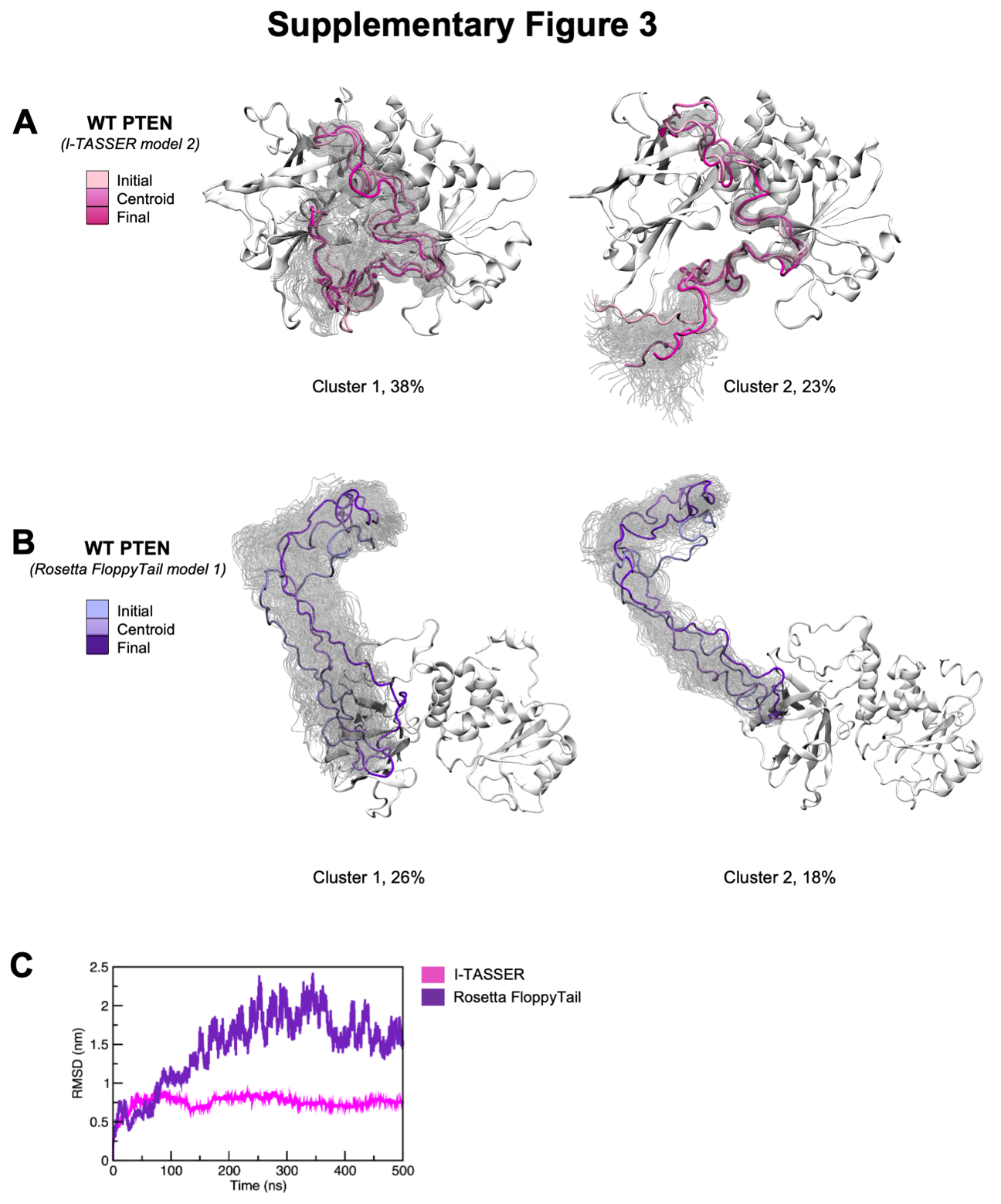

**Figure S4**. Conformational ensemble structures from top two representative centroid clusters of PTEN CTT for (**A**) I-TASSER, WT PTEN second lowest energy model and (**B**) Rosetta FloppyTail, WT PTEN lowest energy model. The backbone RMSD time profiles from the initial structure for each system (**C**). Initial, centroid, and final conformations are colored as a gradient in *magenta* and *violet* for I-TASSER and Rosetta FloppyTail MD simulation replicas, respectively. All other structures in each ensemble are indicated in translucent *gray*. Clusters are from the top two dominant conformational ensembles of I-TASSER (model 2) and Rosetta FloppyTail (model 1), with populations at 38% and 26%, respectively for cluster 1, and at 23% and 18%, respectively for cluster 2.

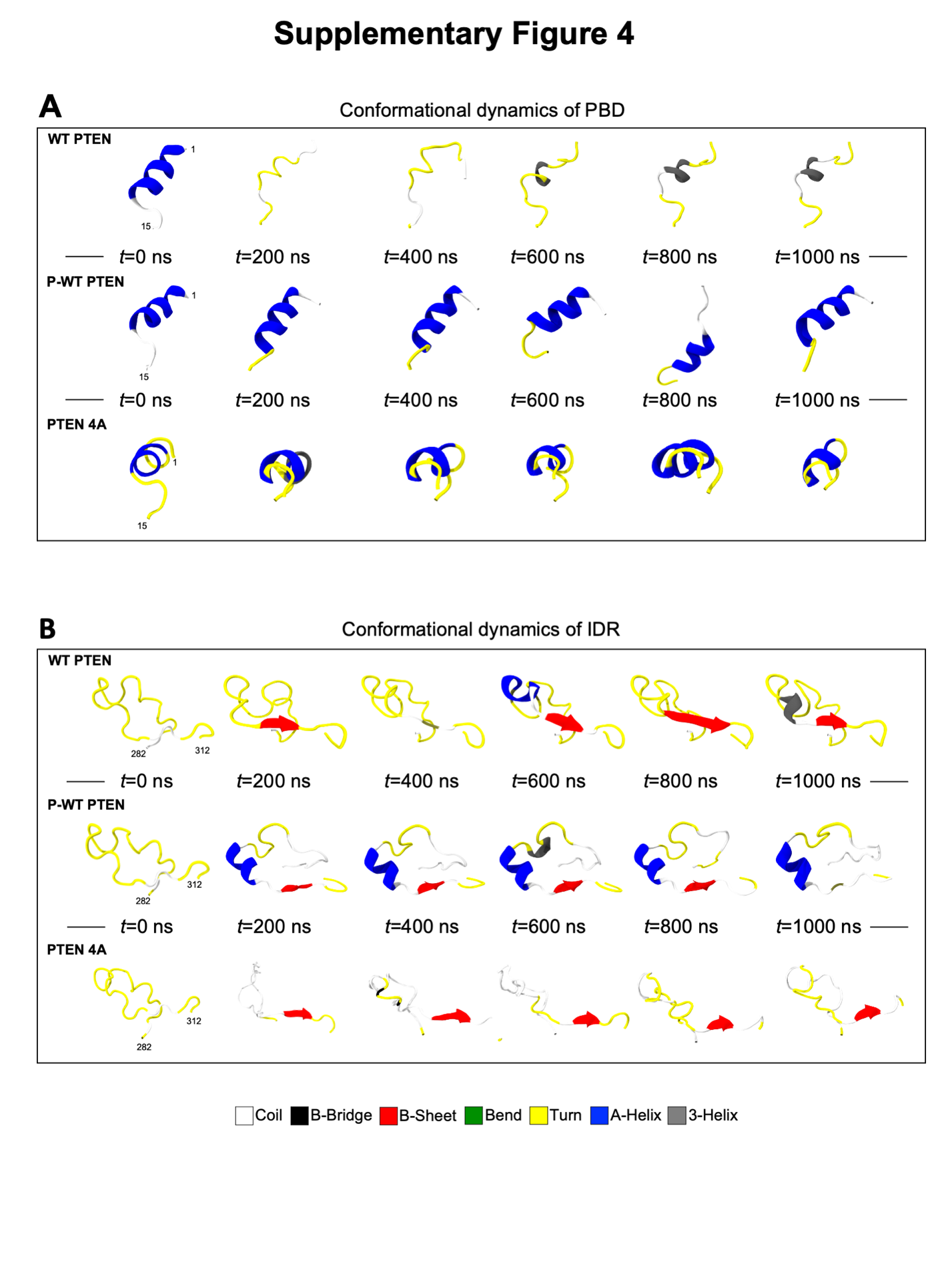

**Figure S5**. Conformational dynamics of PBD and IDR regions. Time-evolution of conformational dynamics of (**A)** PBD (residues 1-15) and (**B)** IDR regions (residues 282-312) over the course of entire MD simulations for WT PTEN (*top* panels), P-WT PTEN (*middle* panels), and PTEN 4A (*bottom* panels), respectively. Descriptions of secondary structure elements are indicated in key.

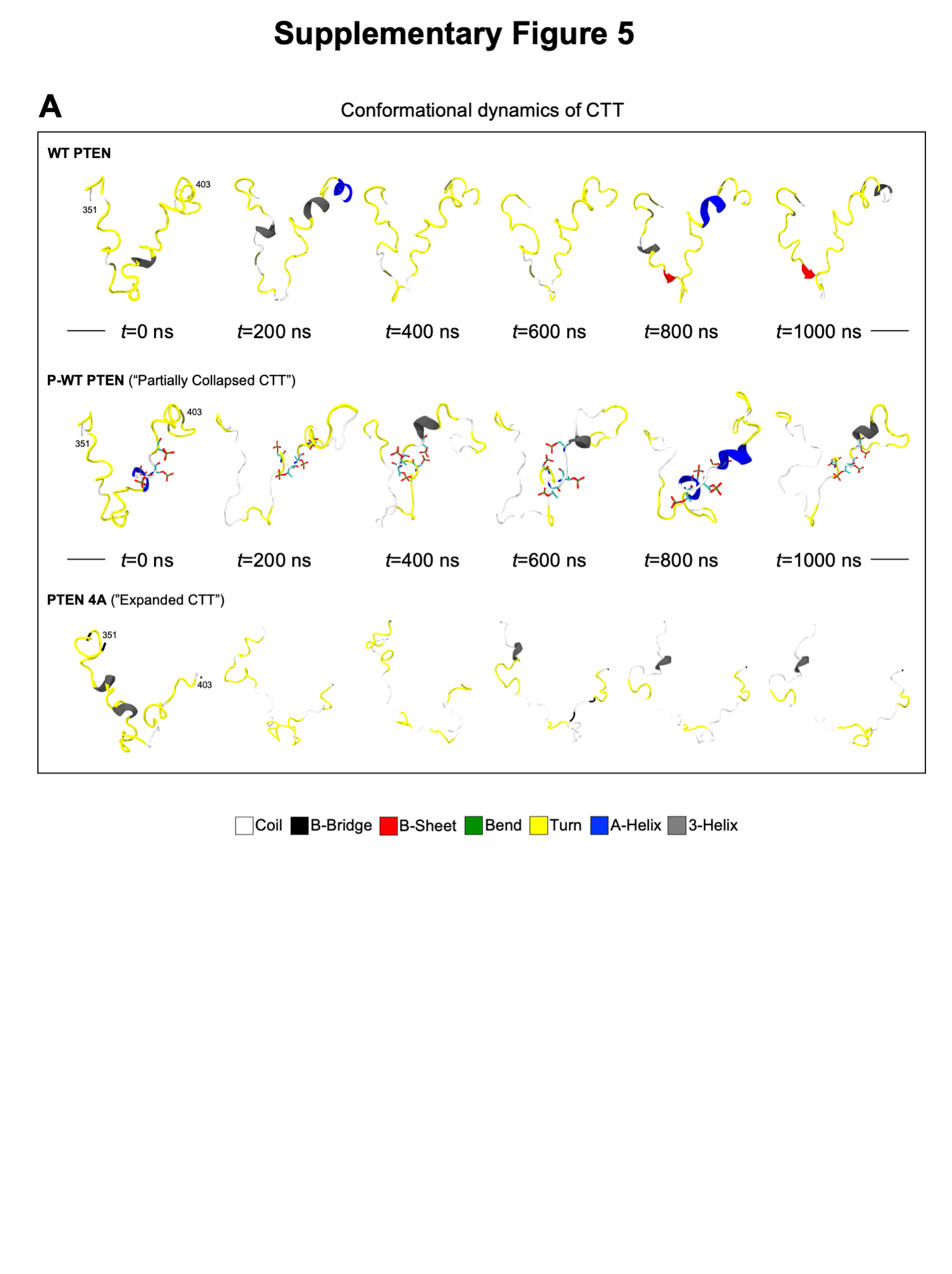

**Figure S6**. Conformational dynamics of CTT region (residues 351-403). Time-evolution of conformational dynamics of (**A)** CTT region over the course of entire MD simulations for WT PTEN (*top* panels), P-WT PTEN (*middle* panels), and PTEN 4A (*bottom* panels), respectively. Descriptions of secondary structure elements are indicated in key.

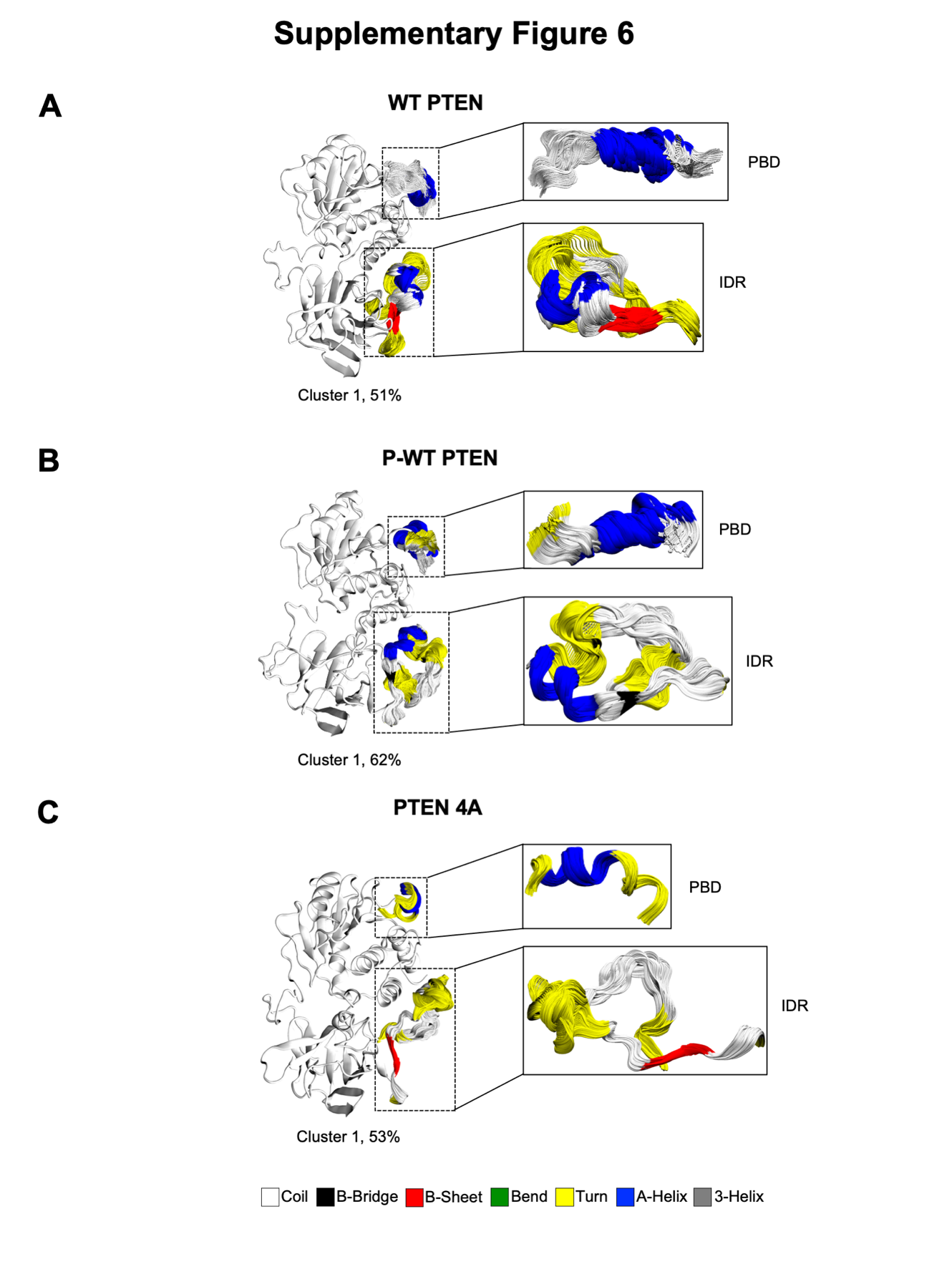

**Figure S7**. Conformational ensemble structures from MD simulation replicas of IDR and CTT regions. To further broadly sample the conformational space, the centroid structures from each most populated cluster from hierarchical cluster analyses served as initial structures of new MD simulations. These clusters are from the most dominant cluster ensemble from subsequent hierarchical cluster analyses of (**A**) WT PTEN, (**B**) P-WT PTEN, and (**C**) PTEN 4A, with populations at 51%, 62%, and 53% respectively. Descriptions of secondary structure elements are indicated in the key.

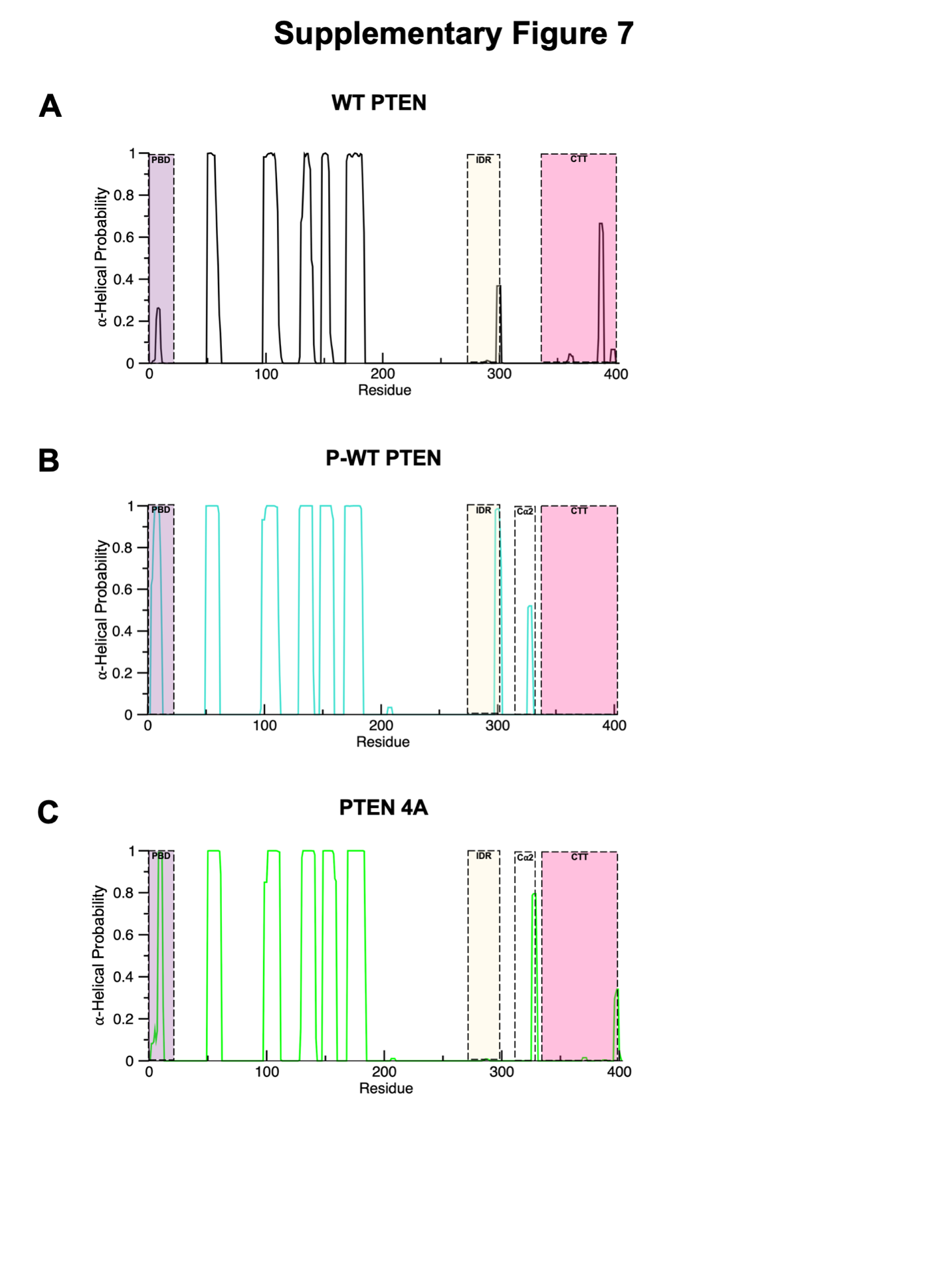

**Figure S8**. Calculation of α-helical probability for (**A**) WT PTEN (*black*), (**B**) P-WT PTEN (*cyan*), and (**C**) PTEN 4A (*green*). PBD (*light purple*), IDR (*light tan*), Cα2 (*white*) and CTT (*light pink*) regions are indicated accordingly.

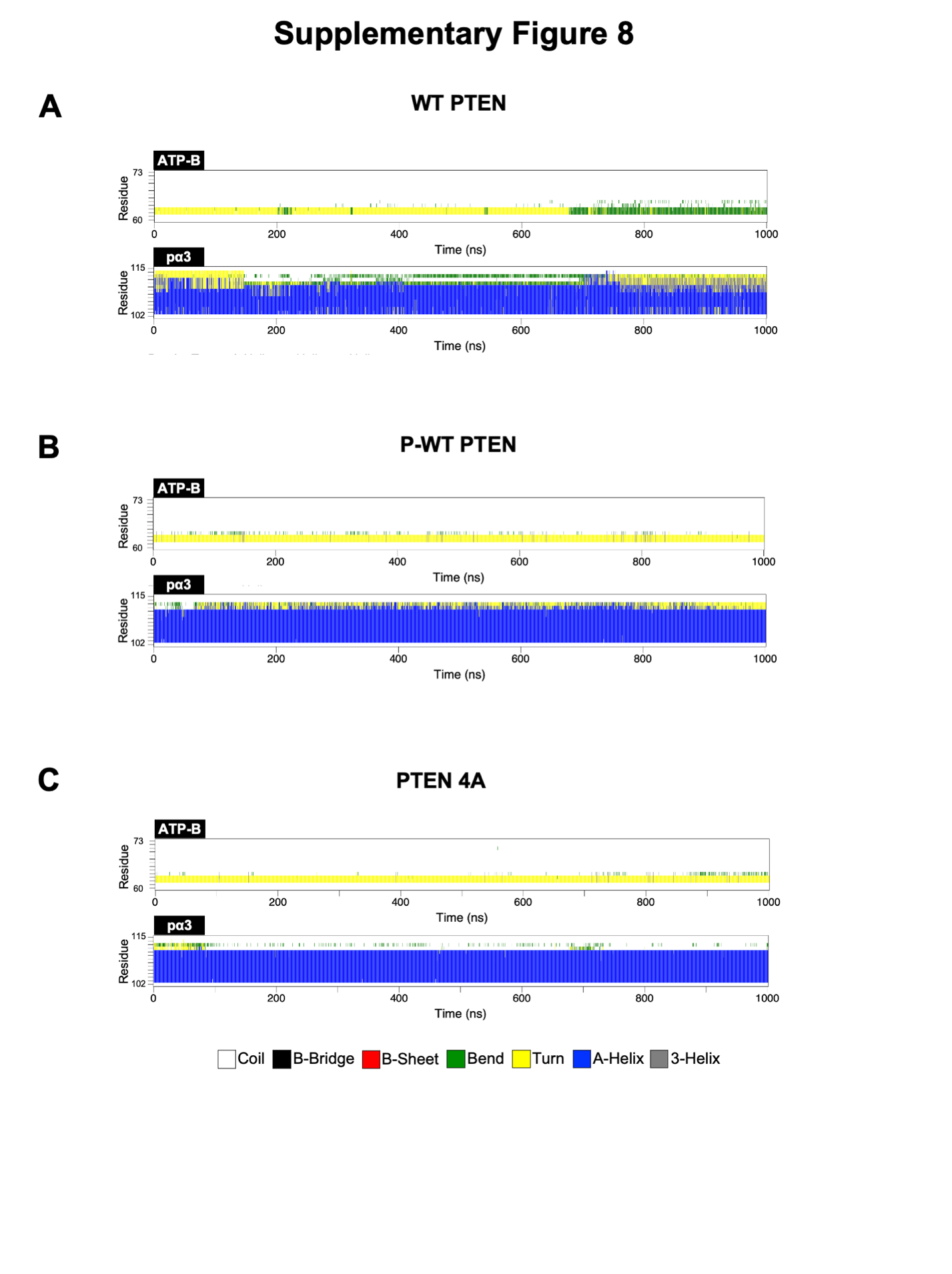

**Figure S9**. Secondary Structure analysis of ATP-B binding motif and pα3 helix for (**A**) WT PTEN, (**B**) P-WT PTEN, and (**C**) PTEN 4A. CTT phosphorylation enhances α-helical formation in pα3 helix of P-WT PTEN. Descriptions of secondary structure elements are indicated in key.

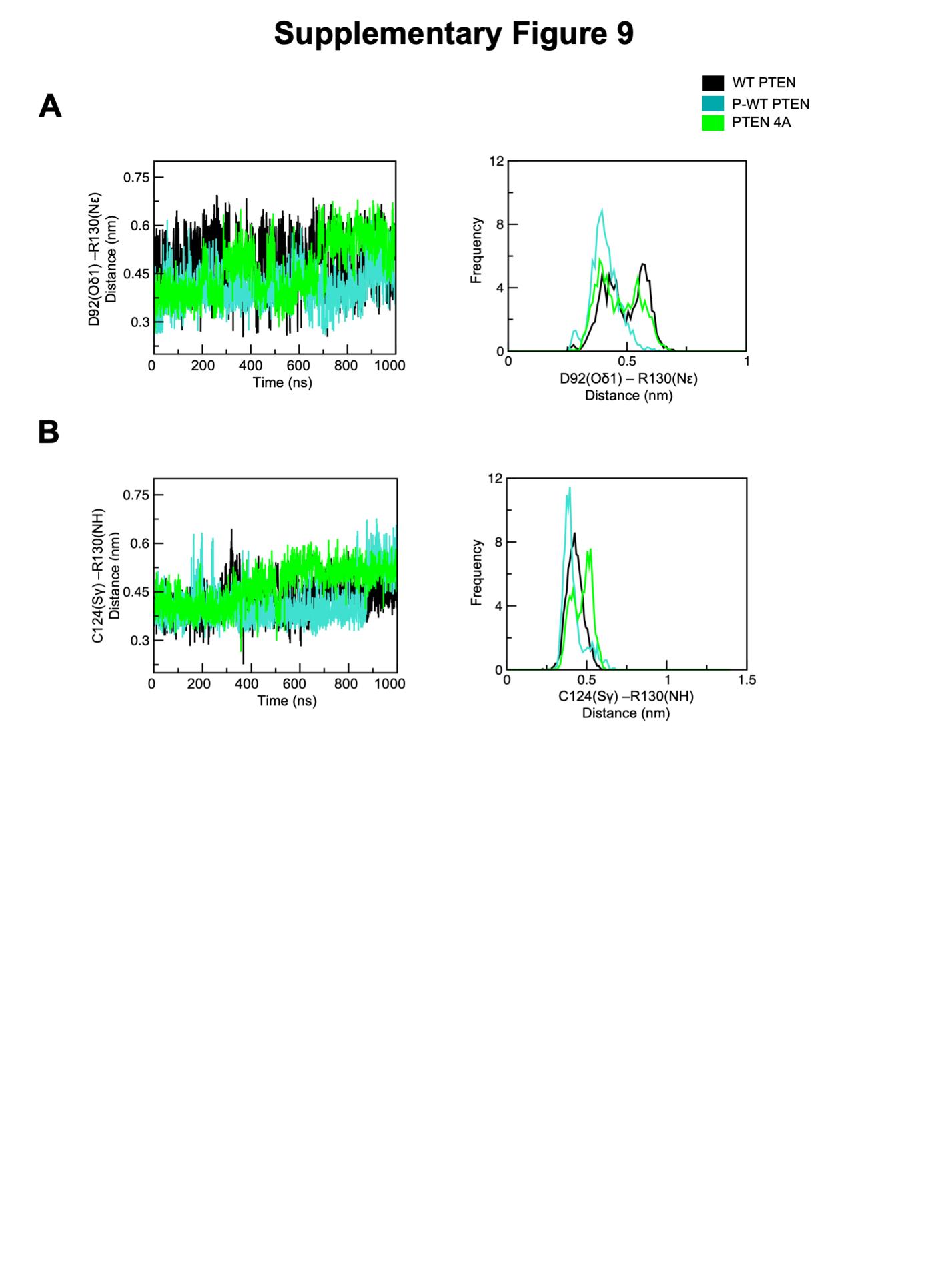

**Figure S10**. Distance interaction between catalytic residues D92-R130 and C124-R130. (**A)** Salt-bridge distance interaction calculated between Oδ1 atom of D92 and Nε atom of R130 and (**B)** Hydrogen bond distance interaction calculated between Sγ atom of C124 and NH atom of R130 for WT PTEN (*black*), P-WT PTEN (*cyan*) and PTEN 4A (*green*) (*left* panel) and frequency of interaction distances (*right* panel).

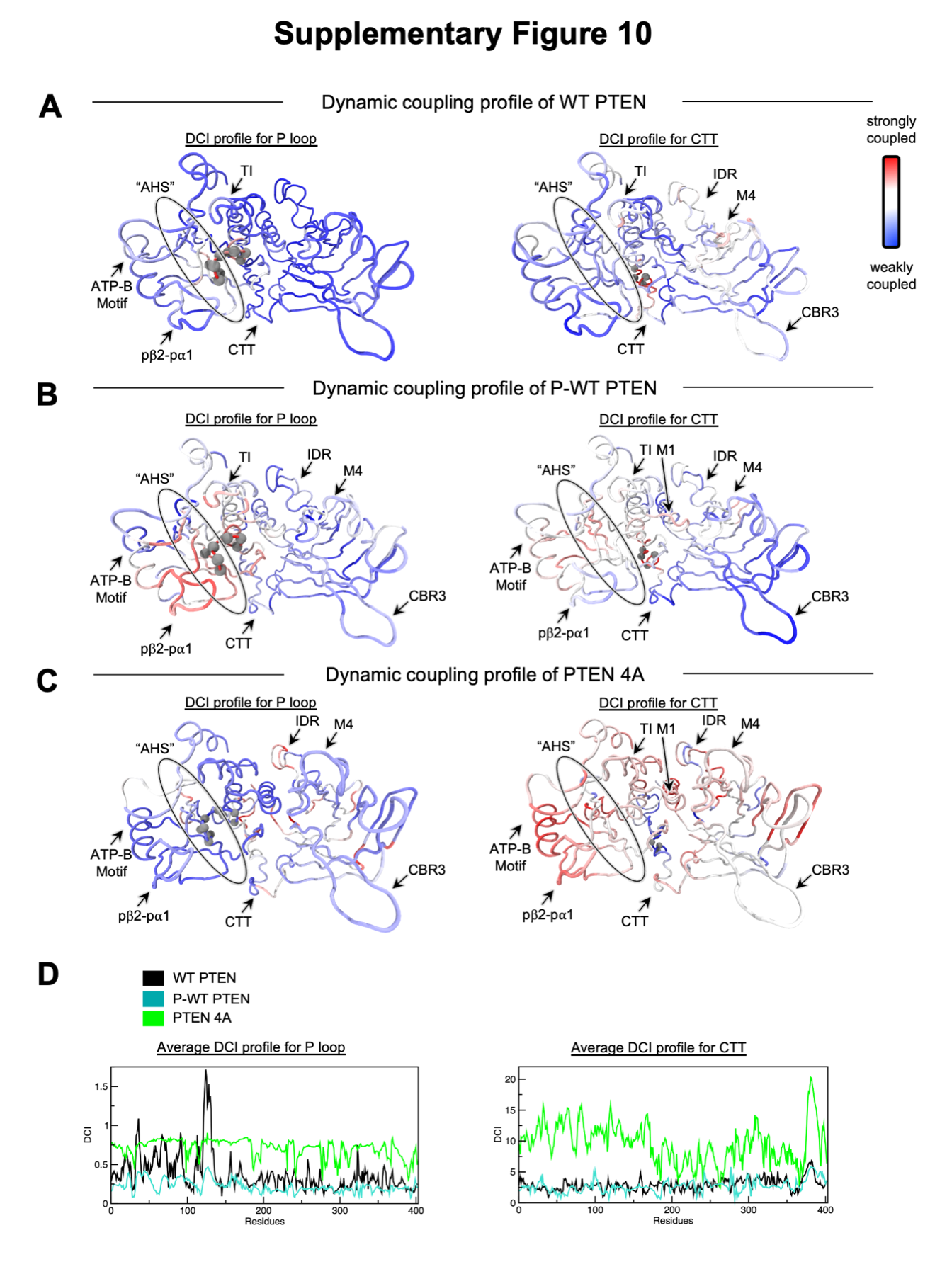

**Figure S11**. Comparison of the coupling of the P loop and CTT regions using dynamic coupling index (DCI). DCI profile differences visualized on the three-dimensional structure for (**A)** WT PTEN P loop (*left* panel) and CTT (*right* panel), (**B)** P-WT PTEN P loop (*left* panel) and CTT (*right* panel), and (**C)** PTEN 4A P loop (*left* panel) and CTT (*right* panel), where *red* represents sites highly coupled to either P loop or CTT (*grey* spheres) and *blue* as sites with no significant coupling. (**D)** Average DCI profile for WT PTEN (*black*), P-WT PTEN (*blue*), and PTEN 4A (*green*).

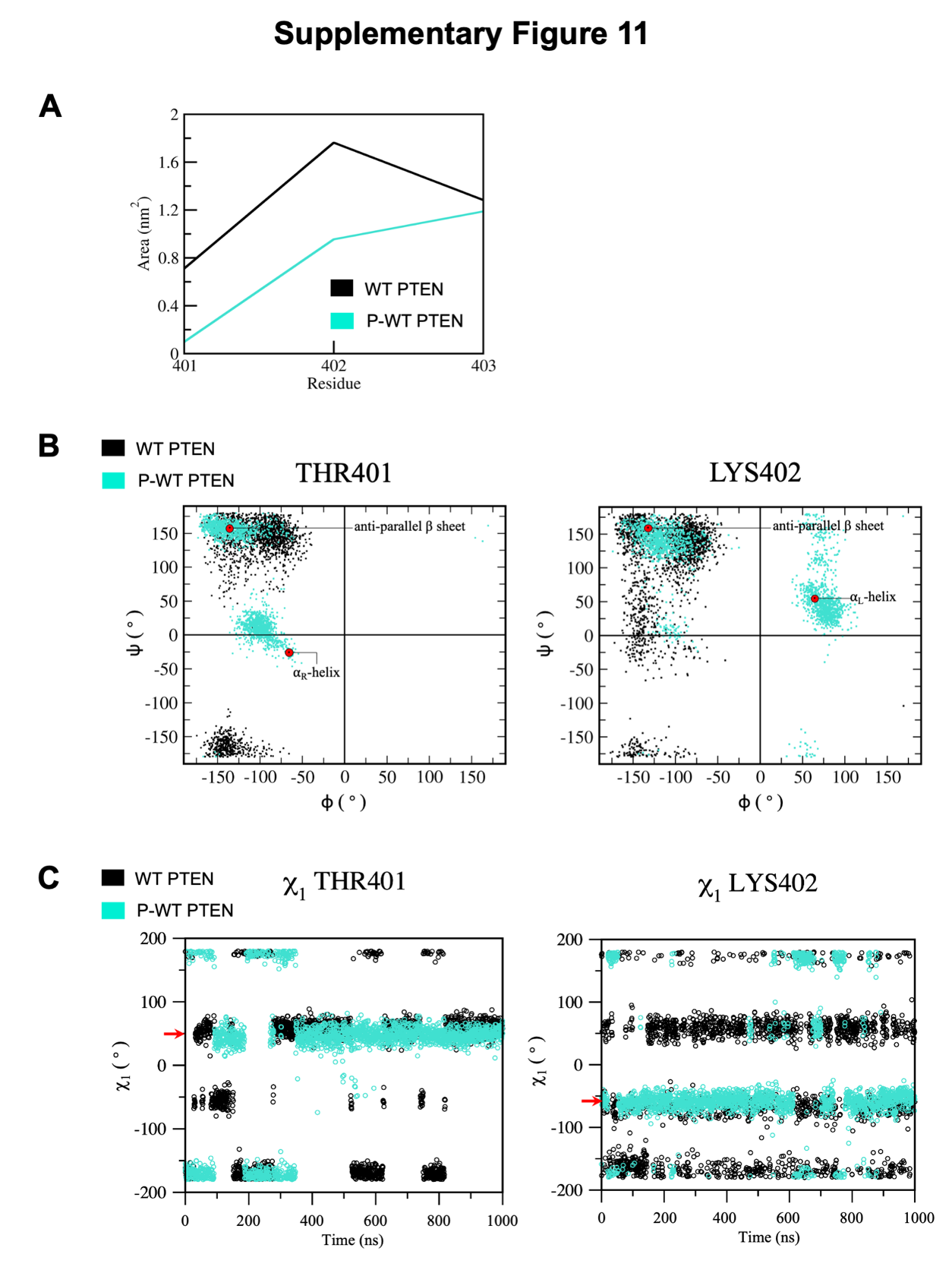

**Figure S12**. Average SASA per residue and comparative analysis of χ_1_ sidechain torsion angle, and Ramachandran plots for PDZ binding motif. (**A)** Average SASA of residues within PDZ binding motif (401TKV403) for WT PTEN (*black*) and P-WT PTEN (*cyan*). (**B)** χ_1_ sidechain torsion angle of PDZ binding motif for WT PTEN (*black*) and P-WT PTEN (*cyan*). *Red* arrows indicate most populated state for P-WT PTEN for T401 (50°) and K402 (-50°). (**C)** Ramachandran plots for WT PTEN (*black*) and for P-WT PTEN (*cyan*). P-WT PTEN shows the phi (φ)-psi (ψ) torsion angles populate an ⍺_R_-helix for T401 and an ⍺_L_-helix for K402 as indicated by the *red* circle.

**Supplementary Table**

**Table S1**. Comparison of the full-length *in silico* PTEN structure (I-TASSER) simulation systems

|  | System | Number of atoms | Simulation time (ns) |
| --- | --- | --- | --- |
| 1 | WT PTEN | 74,155 | 1 x 1,000 |
| 2 | P-WT PTEN | 74,142 | 1 x 1,000 |
| 3 | PTEN 4A | 74,130 | 1 x 1,000 |

**Table S2**. Reproducibility^a^ of the full-length *in silico* PTEN structure (I-TASSER) simulations systems

|  | System | Number of atoms | Simulation time (ns) |
| --- | --- | --- | --- |
| 1 | WT PTEN | 74,155 | 1 x 1,000 |
| 2 | P-WT PTEN | 74,142 | 1 x 1,000 |
| 3 | PTEN 4A | 74,130 | 1 x 1,000 |

^a^Each simulation was initiated from a different seed than initial simulations in **Table S1**.

**Table S3**. Sampling replicas^a^ from different initial full-length *in silico* PTEN structures

|  | System | Number of atoms | Simulation time (ns) |
| --- | --- | --- | --- |
| 1 | WT PTEN  (centroid structure at 473 ns) | 90,097 | 1 x 500 |
| 2 | P-WT PTEN  (centroid structure at 337 ns) | 79,779 | 1 x 500 |
| 3 | PTEN 4A  (centroid structure at 758 ns) | 88,833 | 1 x 500 |

^a^Each simulation was initiated from centroid structure from hierarchical clustering from simulations in **Table S1**.

**Table S4**. Sampling replicas from different full-length *in silico* PTEN models

|  | System | Number of atoms | Simulation time (ns) |
| --- | --- | --- | --- |
| 1 | WT PTEN  (second lowest energy I-TASSER model) | 80,248 | 1 x 500 |
| 2 | WT PTEN  (lowest energy Rosetta FloppyTail model) | 90,316 | 1 x 500 |
