## Supplemental Script for "Structural and Dynamic Effects of PTEN C-terminal Tail Phosphorylation"

*Scripts and commands for Rosetta FloppyTail structure generation*

1. Relaxing initial structures before Rosetta modeling

Rosetta command:

<path_to_Rosetta_directory>/main/source/bin/relax.default.linuxgccrelease -s input.pdb -out:suffix _relaxed @general_relax_flags

Script 1: general_flag_flags

general_relax_flags:

-nstruct 1

-relax:default_repeats 5

-out:path:pdb relax_output/

-out:path:score expected_output/

2. Rosetta FloppyTail protocol

Rosetta command:

<path_to_Rosetta_directory>/main/source/bin/FloppyTail.linuxgccrelease -out:file:silent batchA.o -out:suffix _PBM_batchA @options &

Script 2: options script

#You'll define your own database flag, of course

-database <path_to_Rosetta_directory>/main/database/

#input PDB

-s <input.pbd>

#ex flags give extra rotamers for packing; use_input_sc allows the pre-existing rotamer when packing (useful when paired with sidechain minimization)

-ex1

-ex2

#-use_input_sc

#prevent design, keep the input sequence (use a resfile if you want design)

-packing:repack_only

#minimizer type. I don't know which is best

-run:min_type dfpmin_armijo_nonmonotone

#fragments if you want them; only 3mers are used.

-in:file:frag3 <3mer_fragment_file>

#local

#start of tail

#-FloppyTail:C_root #use when not modeling C-terminal tail

-FloppyTail:flexible_start_resnum 351

#end of tail (assumed to be end of chain)

#chain of tail

-FloppyTail:flexible_chain A

#used for preventing loss of compactness at centroid/fa switch; see documentation

-FloppyTail:short_tail:short_tail_off 0

-FloppyTail:short_tail:short_tail_fraction 1.0

#shear does nothing for extended tails; see documentation

#-FloppyTail:shear_on .33333333333333333333

#turn this one OFF for other uses; activates publication-relevant metrics

#-FloppyTail:publication true

#frequency of full repacking; 10 used for test to fit in refine_cycles

-FloppyTail:refine_repack_cycles 100

#low-end production numbers

-FloppyTail:perturb_cycles 5000

-FloppyTail:refine_cycles 3000

-nstruct 10

#-out:suffix _scratch

3. Scoring FloppyTail structures and finding RMSD of PTEN CTT domain (versus that of initial structure)

Rosetta command:

<path_to_Rosetta_directory>/main/source/bin/score_jd2.linuxgccrelease -database <path_to_Rosetta_directory>/main/database -s <input_file.pdb> -out:file:scorefile score_rms.sc -native <initialstrucfile>.pdb -native_exclude_res 1-350
